# An update to the global Critical Habitat screening layer

**DOI:** 10.1101/2025.04.01.646588

**Authors:** Sebastian Dunnett, Alfred Muge, Alex Ross, Joseph A. Turner, Neil D. Burgess, Matt Jones, Sharon Brooks

## Abstract

The International Finance Corporation (IFC) defines Critical Habitat in Performance Standard 6 (PS6) as high biodiversity value areas requiring net biodiversity gain for projects. We present an updated global screening layer of Critical Habitat aligned with IFC’s 2019 guidance. This layer derives from global datasets covering 53 biodiversity features, categorized as ‘Likely’ or ‘Potential’ Critical Habitat based on agreement with IFC criteria and data suitability. Analysis indicates 52.77 million km^2^ (10.4%) and 15.94 million km^2^ (3.2%) of the globe can be considered Likely and Potential Critical Habitat respectively, with the remaining 86.4% not overlapping with assessed biodiversity features. This represents a significant increase over previously identified 5% and 2% for Likely and Potential Critical Habitat. Likely Critical Habitat was dominated by Important Bird and Biodiversity Areas, Intact Forest Landscapes, and protected areas; Potential Critical Habitat by Important Marine Mammal Areas and ranges of IUCN Vulnerable species. Our results can help businesses prioritize impact avoidance and identify opportunities by screening potential development sites for biodiversity features.

## Background & Summary

More than US$ 60 trillion of infrastructure spending is likely required to meet 2040 societal goals, with an additional 1.2 million km^2^ urbanised land by 2030, an additional 3-4.7 million km of roads by 2050^1,2,3^. During the relatively modest infrastucture expansion of the late twentieth and early twenty-first centuries, biodiversity has continued to decline precipitously, with governments collectively failing every single one of the Aichi Biodiversity Targets set under the United Nations (UN) Convention on Biological Diversity (CBD) in 2011. Much of the potential expansion of infrastructure is predicted to occur in some of the world’s highest integrity ecosystems ^4,5,6,7^.

In 2012, the International Finance Corporation (IFC; a member of the World Bank Group) revised their *Performance Standard 6: Biodiversity Conservation and Sustainable Management of Living Natural Resources* (hereafter PS6), one of eight Performance Standards any client of the organisation must meet throughout the life of an investment^8^. PS6 draws heavily on fundamental conservation principles such as protected areas and threatened species ^9,10,11^. The standard is seen as the benchmark for sustainable investment with 130 financial institutions following its social and environmental classification process as signatories to the Equator Principles ^12^. Businesses can prioritise impact avoidance and direct corporate nature action with Critical Habitat screening ^13^, even more important when, for example, half of marine Critical Habitat identified in 2015 is not formally protected and would not be flagged when only considering legally designated sites ^14^.

Critical Habitat is defined by IFC using the following criteria:

1. Critically Endangered or Endangered species
2. Endemic and/or restricted-range species
3. Globally significant concentration of migratory or congregatory species
4. Highly threatened and/or unique ecosystems
5. Key evolutionary processes

Previous work by Martin *et al*. ^15^ and Brauneder *et al*. ^16^ produced global 1 km^2^ IFC Critical Habitat screening layers for the marine and terrestrial realms respectively. These studies identified global biodiversity feature data with sufficient resolution and alignment with IFC PS6 criteria to produce global screening layers for Likely and Potential Critical Habitat. Martin *et al*. ^15^ classified 1.6% of the ocean as Likely Critical Habitat and 2.1% as Potential Critical Habitat. Brauneder *et al*. ^16^ identified 10% of the terrestrial surface as Likely Critical Habitat and 5% as Potential Critical Habitat. The screening layers, as well as the amalgamated marine plus terrestrial Critical Habitat layer, are made available through the UN Environment Programme World Conservation Monitoring Centre’s [data portal](https://data-gis.unep-wcmc.org/portal/home/) as well as the [UN Biodiversity Lab](https://unbiodiversitylab.org/en/). The layers have been used in a number of subsequent studies, including analysis of the impact of PS6 on biodiversity offset implementation ^17^. Many of the studies focus on the threat to biodiversity of China’s Belt and Road Initiative, a programme to link sixty-five countries with a network of transport and energy infrastructure. Researchers have used the global Critical Habitat screening layer to estimate that 50% of loans intersect with Critical Habitat^1 18^. Full 49 Chinese-funded dams have a total of 149 km^2^ Critical Habitat in close proximity (i.e. within 1 km) and dams funded by multilateral development banks (MDBs) have much lower average areas of Critical Habitat per area of dam at risk relative to Chinese-funded dams ^19^. Researchers have also identified 12,000 km^2^ of Critical Habitat within 1 km of the Belt and Road Initiative’s linear infrastructure ^20^. In a decarbonising world, 20% of mining locations in Africa trigger a Critical Habitat classification for something other than Great Ape habitat ^21^.

In June 2019, IFC updated the 2012 PS6 Guidance Note that had accompanied the initial release ^22^. The update significantly changed the Critical Habitat criteria to better align the standard with the then newly developed *Global Standard for the Identification of Key Biodiversity Areas* (KBAs), Table 1. This meant that the original Critical Habitat global screening layers were now out of sync with the latest criteria. At the same time, many of the datasets that underpinned the original analyses have been updated multiple times since (e.g. the the World Database on Protected Areas – WDPA – is updated monthly). Finally, many eligible datasets have been produced in the years since that meet the completeness and alignment criteria of the layer.

**Table 1.** Comparison of thresholds for triggering Critical Habitat in the update Guidance Note. Implications are based on changes to the lowest threshold for Critical Habitat (i.e. Tier 2 in 2012). Adapted from UNEP-WCMC^23^.

| Criteria | GN 2012 | GN 2019 | Implications |
| --- | --- | --- | --- |
| Critically Endangered or Endangered species | Tier 1: 10%<br>Tier 2: >0% | 0.5% | Small increase in threshold<br>Reduction in overall Likely or Potential Critical Habitat |
| Endemic and/or restricted-range species | Tier 1: 95%<br>Tier 2: 1% | 10% | Threshold significantly higher<br>Reduction in overall Likely or Potential Critical Habitat |
| Globally significant concentrations of migratory or congregatory species | Tier 1: 95%<br>Tier 2: 1% | 1% | Threshold unchanged<br>No change to overall Likely or Potential Critical Habitat |
| Highly threatened and/or unique ecosystems | Qualitative<br>No thresholds | 5% | New threshold |
| Key evolutionary processes | Qualitative<br>No threshold | Qualitative<br>No threshold | Unchanged |

For businesses to manage their biodiversity impacts in areas of high biodiversity value, it is essential to provide an updated Critical Habitat layer that fully aligns with the 2019 update of PS6. In this Data Descriptor, we describe a new, updated global Critical Habitat screening layer. The new data incorporates updated datasets, as well as new sources identified through a comprehensive search for any that could meet the eligibility criteria. Most importantly, we describe and make available a methodology whereby the layer can be updated as and when new data are found or updated.

## Methods

### Critical Habitat criteria

IFC define Critical Habitat according to five criteria (Table 1) with associated thresholds for Criteria 1-4. The 2019 update to the accompanying Guidance Note made a number of changes to the classification of Critical Habitat of relevance to any updated methodology:

- **Changes to thresholds**: thresholds have been updated to better align with the Global KBA Standard ^24^. Criteria 1, 3 and 4 now directly align with the standard, and Criteria 1-3 have streamlined thresholds that largely sit between the previous Tier 1 and Tier 2 thresholds. Quantitative thresholds have been added for Criterion 4, based on the developing IUCN Red List of Ecosystems ^25^.
- **Specified exceptional circumstances**: IFC now specify three circumstances that trigger special considerations.
  – Great Apes: the presence of Great Ape species is likely to trigger Critical Habitat regardless of thresholds. Where present, clients are expected to inform both IFC and the IUCN Species Survival Commission’s Primate Specialist Group.
  – Unapprovable sites:
    * World Heritage Sites (Natural or Mixed).
    * Alliance for Zero Extinction sites.
- **Addition of IUCN Vulnerable species**: IFC added a threshold for Criteria 1 that now explicitly includes species classified as Vulnerable (VU) by the IUCN Red List of Threatened Species (hereafter IUCN Red List) where:
  – Globally important populations are present.
  – The loss of the population would lead to the species being upgraded to Endangered (EN) or Critically Endangered (CR).
  – The population in question would then trigger Criterion 1a (the area supports >=0.5% of the global population and >=5 reproductive units).

### Data screening and classification

Using the updated criteria, we proceeded with data screening and classification as with previous studies ^15,16^. Datasets were identified through expert knowledge and consultation, using the criteria adapted from Martin *et al*. ^15^:

1. Direct relevance to one or more Critical Habitat criteria.
2. Global in extent.
3. Assembled using a standardised protocol.
4. The best available data for the biodiversity feature of interest.
5. Sufficiently high resolution to indicate presence of biodiversity on the ground at scales relevant to business operations.

Datasets that met our criteria were classified as one of three classes: Likely, Potential, and Unclassified. The classification was done by expert judgement, weighing the alignment with PS6 criteria and the likely presence on the ground (Figure 1).

**Figure 1.**
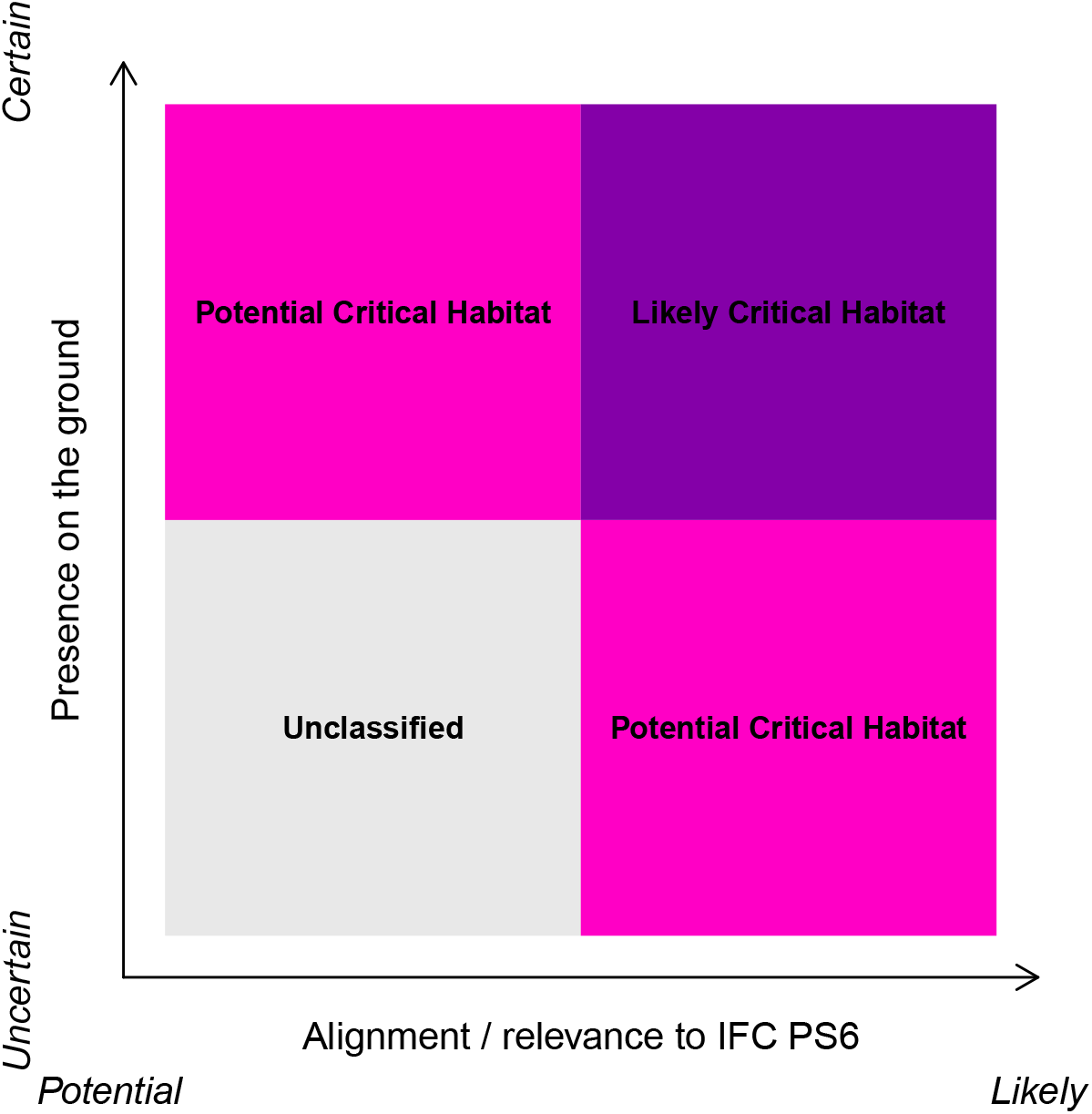
Classification of data as Likely or Potential Critical Habitat is based on the perceived strength of alignment with PS6 criteria and the spatial resolution of the data. Adapted from Brauneder et al. ^16^.

We identified 22 biodiversity feature datasets that are split into 54 separate triggers (Table 2).

**Table 2.**
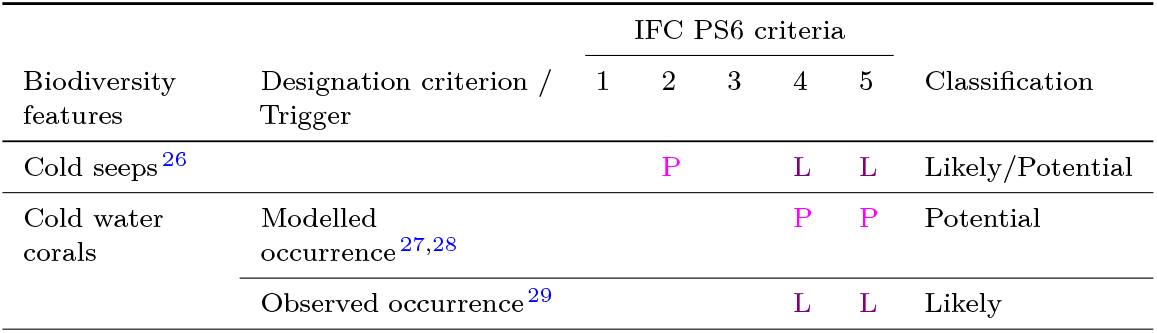
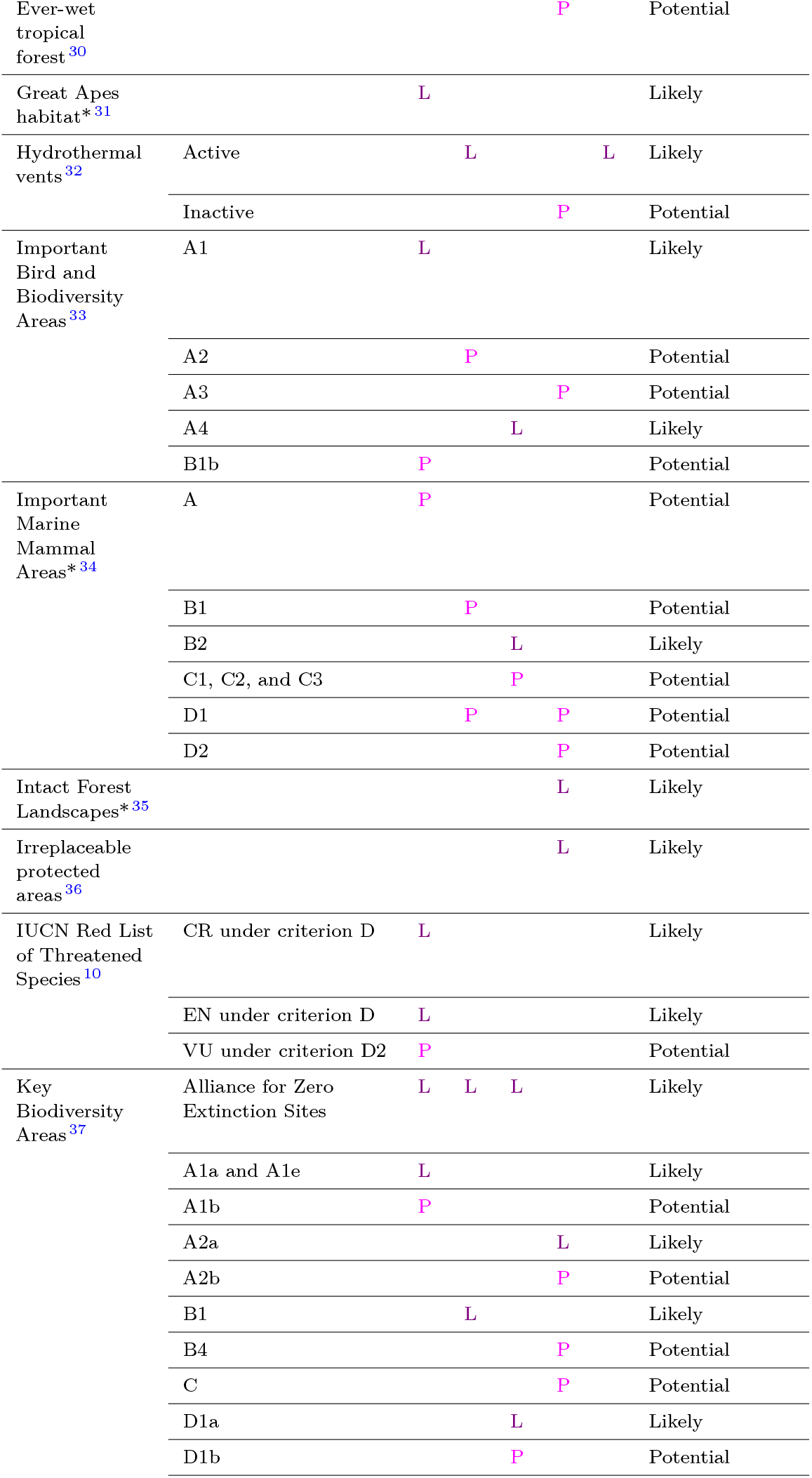
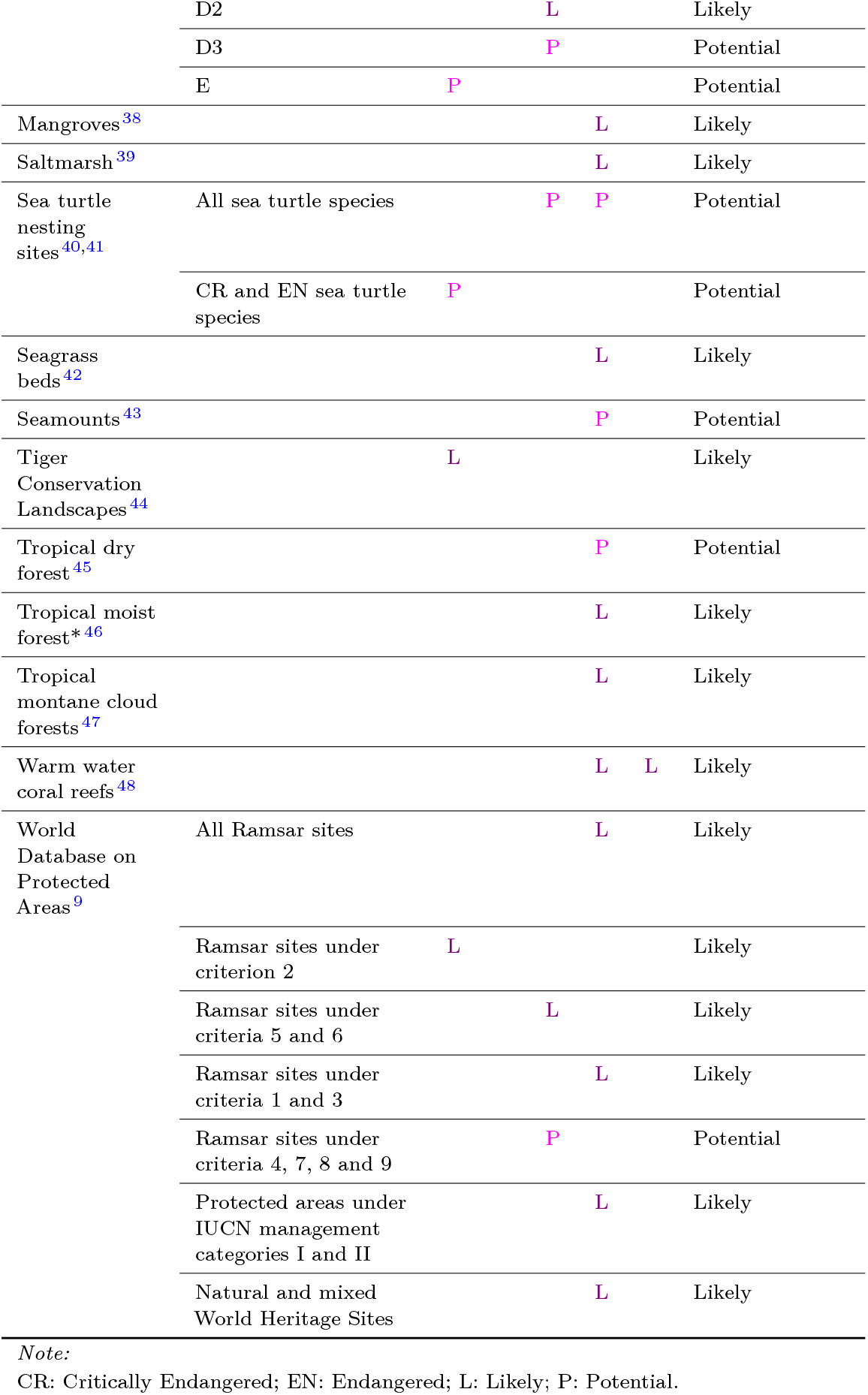
Biodiversity features included in the analysis, their alignment with IFC PS6 Critical Habitat criteria and classification as ‘Likely’ or ‘Potential’ Critical Habitat. *Data newly added in the update.

### Data processing and spatial analysis

Spatial data analysis was conducted in **R**, predominantly using the `terra` and `sf` packages.

For the greatest accuracy, vector data were processed using the `sf` package and `S2` library. This allows spatial processing, e.g. intersections and buffers, using Great Circle distances. Data were prepared for the analysis in three steps: First, data were made valid on the sphere. Second, data were filtered as detailed by (a) their respective guidance, and (b) to produce the data subset required for the analysis. For example, Key Biodiversity Areas (KBAs) must first be filtered by those whose status is confirmed (`KbaStatus==`“`confirmed`”) and then by the criterion/criteria required to create the subset (e.g. sites designated under KBA Criterion B1). Finally, data were unioned to remove duplicates from point data and any overlapping area in polygon data. Not all input data were available in vector format. For raster data, preprocessing varies by source but generally involved aggregating to the correct resolution (30 arcseconds) before reprojecting the data to a template raster in WGS 84.

Data are then converted to binary maps of presence/absence. For vector data, this was done by a process called rasterisation, whereby any grid cell in contact with the vector data assumes the value of the vector data. In keeping with the precautionary nature of the screening layer, we chose to rasterise based on any intersection, not requiring polygon data to cross the midpoint of the grid cell. Point data transferred their values to the grid cells they occupy. For raster data, where necessary (i.e. the data were not already binary), a classification threshold was set. Again, in keeping with the precautionary nature of the data, we set this at 0.5. This meant that any cell with >=50% of the feature was classified as a presence (see Technical Validation for a brief sensitivity analysis of this threshold).

Binary raster data were then combined so that each grid cell’s unique combination of biodiversity features has a unique value. Based on what features comprised the value, the grid cell was categorised hierarchically: Likely > Potential > Unclassified.

## Data Records

We find that 67.79 million km^2^ of the Earth’s surface was classified as Likely or Potential Critical Habitat (Figure 2): 53.81 million km^2^ (10.55%) as Likely Critical Habitat and 13.98 million km^2^ (2.74%) as Potential Critical Habitat. This is a significant increase on the 25.78 million km^2^ (5.05%) and 10.1 million km^2^ (1.98%) previously identified. The remaining 442.3 million km^2^ (86.71%) is “Unclassified” as either known biodiversity features do not align with the IFC definition or because appropriate data that might be used to classify do not exist: Critical Habitat may still occur in these regions.

**Figure 2.**
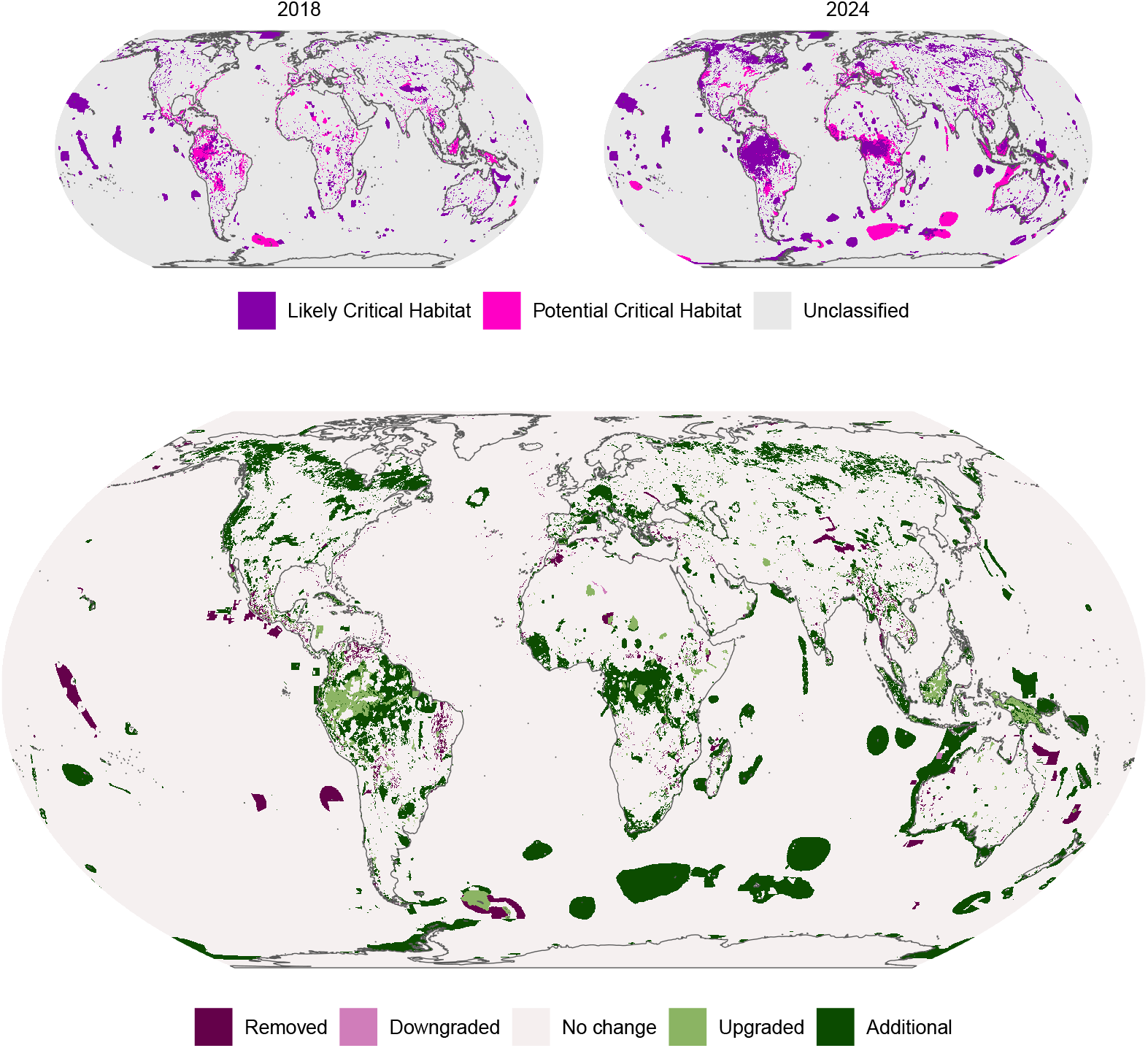
Global screening layer for Critical Habitat. Reprojected to Equal Earth and aggregated to 10 × 10 km for visualisation.

These data are presented in two formats at 30 arcseconds resolution in WGS 84 projection and one in WGS 84 vector format. They represent the best publicly available global screening layer for Critical Habitat:

### Basic Critical Habitat layer

*Basic_Critical_Habitat.tif*

Basic global layer at 30 arcseconds resolution in WGS 84 projection containing three values: 0 (Unclassified), 1 (Potential Critical Habitat), and 10 (Likely Critical Habitat). Made available under a [CC BY licence](https://creativecommons.org/licenses/by/4.0/) at https://doi.org/10.34892/snwv-a025.

### Drill down Critical Habitat layer

*Drill_Down_Critical_Habitat.tif* and *Drill_Down_Critical_Habitat.tif.vat.dbf*

More detailed global layer at 30 arcseconds resolution in WGS 84 projection. The spatial grid (*Drill_Down_Critical_Habitat.tif*) works in conjunction with the raster attribute table (RAT: *Drill_Down_Critical_Habitat.tif.vat.dbf*) to detail what biodiversity features trigger any cell’s Critical Habitat classification. Values in the `TIFF` file correspond to unique identifiers in the RAT that reveal the underlying data (Table 3). Made available under a CC BY-NC licence at https://doi.org/10.34892/d3xm-qm60.

**Table 3.**
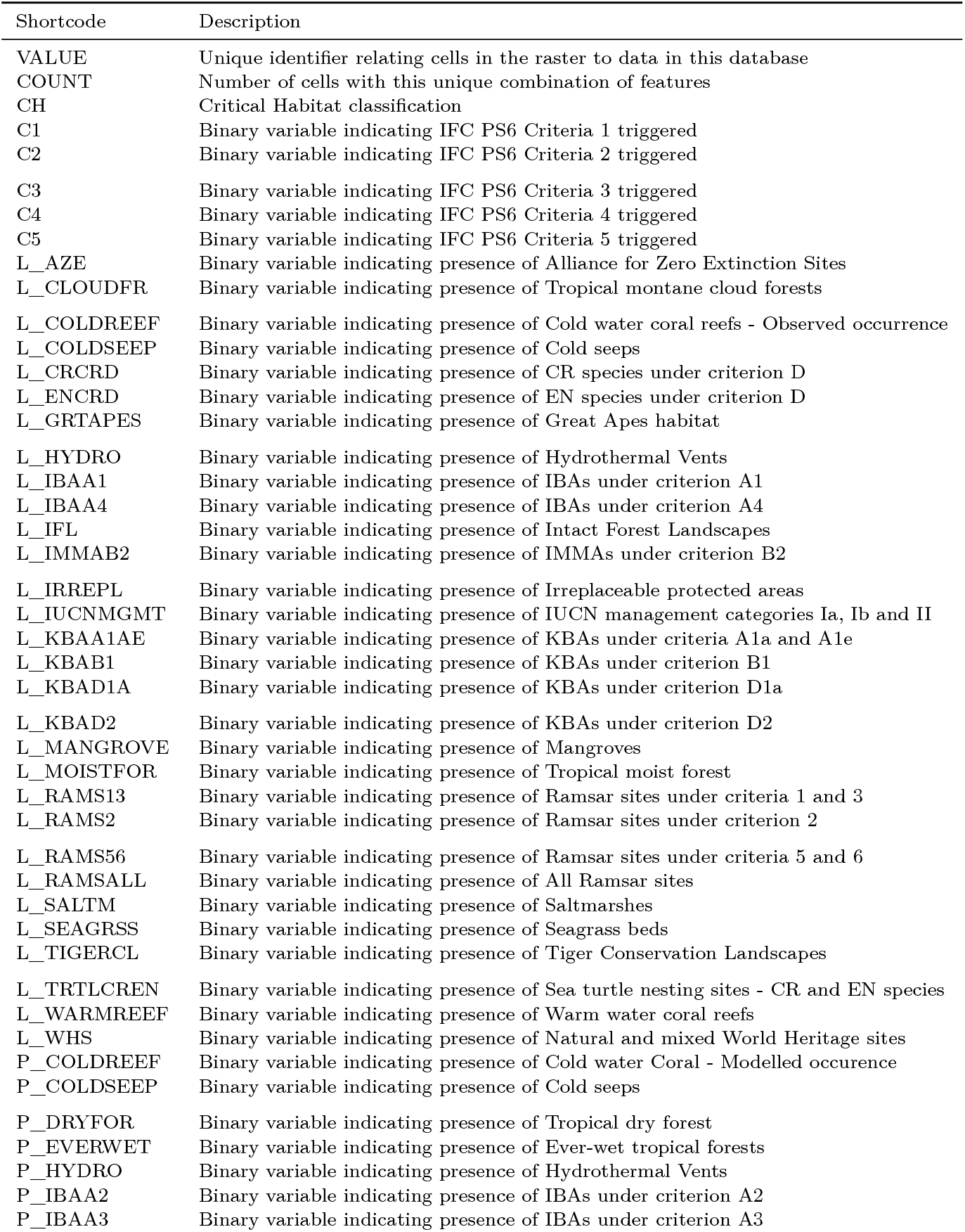
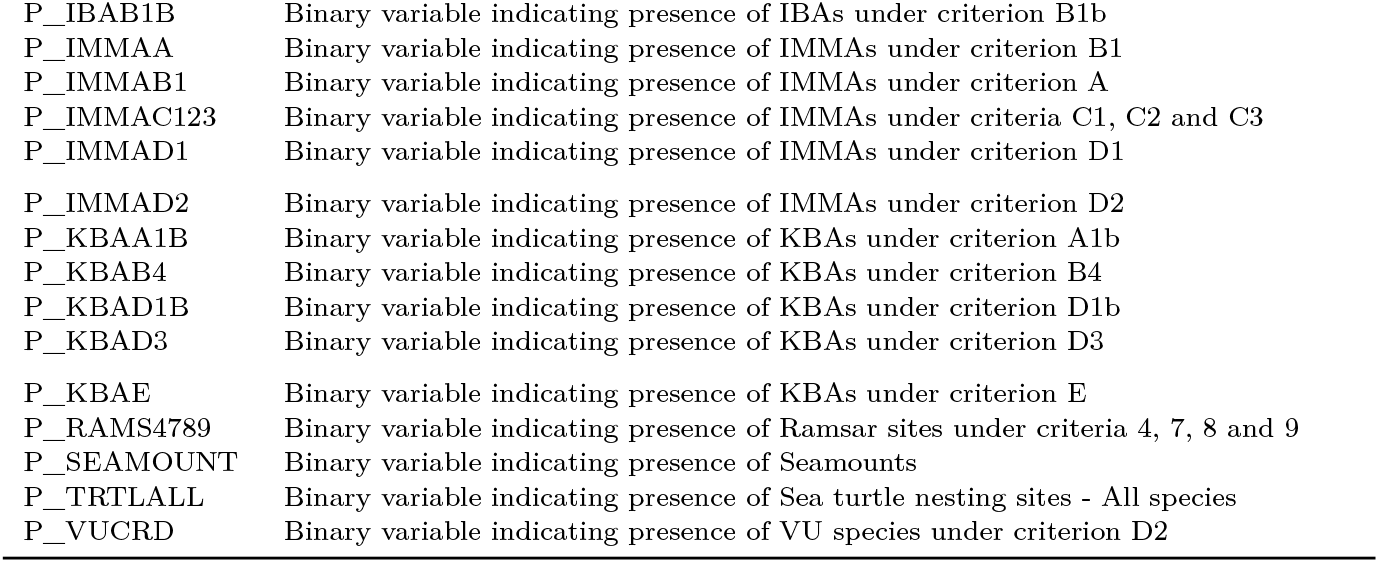
Codes for biodiversity features in the raster attribute table.

### Drill down Critical Habitat layer - polygons

*Drill_Down_Critical_Habitat_Polygons.gpkg*

This polygonised version of the drill down raster layer allows for more concise information retention (Table 4). The data draw boundaries around the cells of each unique feature combination. For ease, and to limit the size of the file, we excluded Unclassified polygons. There are 20,410 polygons in the GeoPackage. Made available under a CC BY-NC licence.

**Table 4.** Variable names and descriptions in the polygonised GeoPackage layer.

| Name | Description |
| --- | --- |
| CH | Critical Habitat designation: Likely, Potential, or Unclassified. |
| CRITERIA | IFC PS6 Criteria triggered. |
| ALL_FEATURES | All biodiversity features found in polygon. |
| C1 | All biodiversity features triggering IFC PS6 Criteria 1. |
| C2 | All biodiversity features triggering IFC PS6 Criteria 2. |
| C3 | All biodiversity features triggering IFC PS6 Criteria 3. |
| C4 | All biodiversity features triggering IFC PS6 Criteria 4. |
| C5 | All biodiversity features triggering IFC PS6 Criteria 5. |

### Technical validation

#### Coverage

Both categories of Critical Habitat increased their absolute areas relative to the first global screening layer. Likely Critical Habitat maintained a largely similar split proportionally across domains (Table 5) but Potential Critical Habitat coverage increased in Areas Beyond National Jurisdiction (ABNJ) relative to land and Exclusive Economic Zones (EEZ). Overall, the updated screening layer increased coverage in all domains, with a large increase from 1.10% to 3.70% in ABNJ, 8.02% to 13.85% in EEZ and 15.09% to 27.29% on land.

**Table 5.** Areas and percentage coverage of each Critical Habitat category in 2018 and 2024 across realms. Percentages indicate splits within Critical Habitat categories.

| Domain | 2018 |  |  |  |  |  | 2024 |  |  |  |  |  |
| --- | --- | --- | --- | --- | --- | --- | --- | --- | --- | --- | --- | --- |
|  | Likely |  | Potential |  | Unclassified |  | Likely |  | Potential |  | Unclassified |  |
| ABNJ | 2.4 | 9.2% | 0.052 | 0.6% | 220 | 46.4% | 5.1 | 9.4% | 3.2 | 22.6% | 214 | 48.4% |
| EEZ | 8.8 | 34.2% | 2.472 | 24.4% | 129 | 27.2% | 14.5 | 26.8% | 5.0 | 35.8% | 121 | 27.4% |
| Land | 14.6 | 56.6% | 7.578 | 75.0% | 125 | 26.4% | 34.3 | 63.8% | 5.8 | 41.4% | 107 | 24.2% |

*Note:*
Areas in km<sup>2</sup> x 10<sup>6</sup>. ABNJ: Areas Beyond National Jurisdiction; EEZ: Economic Exclusion Zone.

Table 6 sets out the coverage of the Critical Habitat screening layer across the five criteria. Criteria 1, 3, and 4 dominate coverage. As identified previously ^15,16^, data are still either largely unavailable or do not meet Criterion 5, key evolutionary processes, and all biodiversity features triggering it remain in the marine realm. The ongoing development and improved coverage of the IUCN Red List of Ecosystems will likely serve to harmonise much of the data inputs to Criterion 4 and reduce the percentage contribution of Criterion 4 to Likely and Potential Critical Habitat classifications ^25,49^.

**Table 6.** Areas and percentage cover of each IFC PS6 criteria to Potential and Likely Critical Habitat. Percentages indicate splits within Critical Habitat categories.

| Criteria | Likely |  | Potential |  |
| --- | --- | --- | --- | --- |
| 1. Critically Endangered or Endangered species | 27,896 | 51.8% | 7,956 | 57.0% |
| 2. Endemic and/or restricted-range species | 14,482 | 27.0% | 1,747 | 12.6% |
| 3. Globally significant concentration of migratory or congregatory species | 21,026 | 39.0% | 6,995 | 50.0% |
| 4. Highly threatened and/or unique ecosystems | 38,904 | 72.2% | 4,641 | 33.2% |
| 5. Key evolutionary processes | 535 | 1.0% | 725 | 5.2% |

*Note:*
Areas in km<sup>2</sup> x 10<sup>3</sup>. Percentages add up to >100 due to overlapping areas.

#### Large species ranges

The inclusion of species classified as Vulnerable in the IFC Guidance Note 2019 update, and data updates to the IUCN Red List generally, necessitated the inclusion of large numbers of species ranges. Some of these ranges, e.g. the Australian grey falcon, *Falco hypoleucos*, are exceptionally large. In order to retain precision in the updated screening layer, we conducted a sensitivity analysis of trimming the largest species ranges (range areas 3, 10, 20 and 50 standard deviations above the mean; see Figure 3 below). Trimming species ranges 3 standard deviations above the mean resulted in a 61.2% decrease in coverage for the removal of only 31 species ranges of 3,953. Furthermore, among the species ranges excluded were the northern white rhino, *Ceratotherium simum cottoni*, whose species range spans several Central and Eastern African countries despite comprising only two living individuals, and the ivory-billed woodpecker, *Campephilus principalis*, which is presumed extinct. Species are not expected to be evenly distributed within this range and if it is otherwise important for threatened species it will likely be classified as Critical Habitat via other biodiversity features.

**Figure 3.**
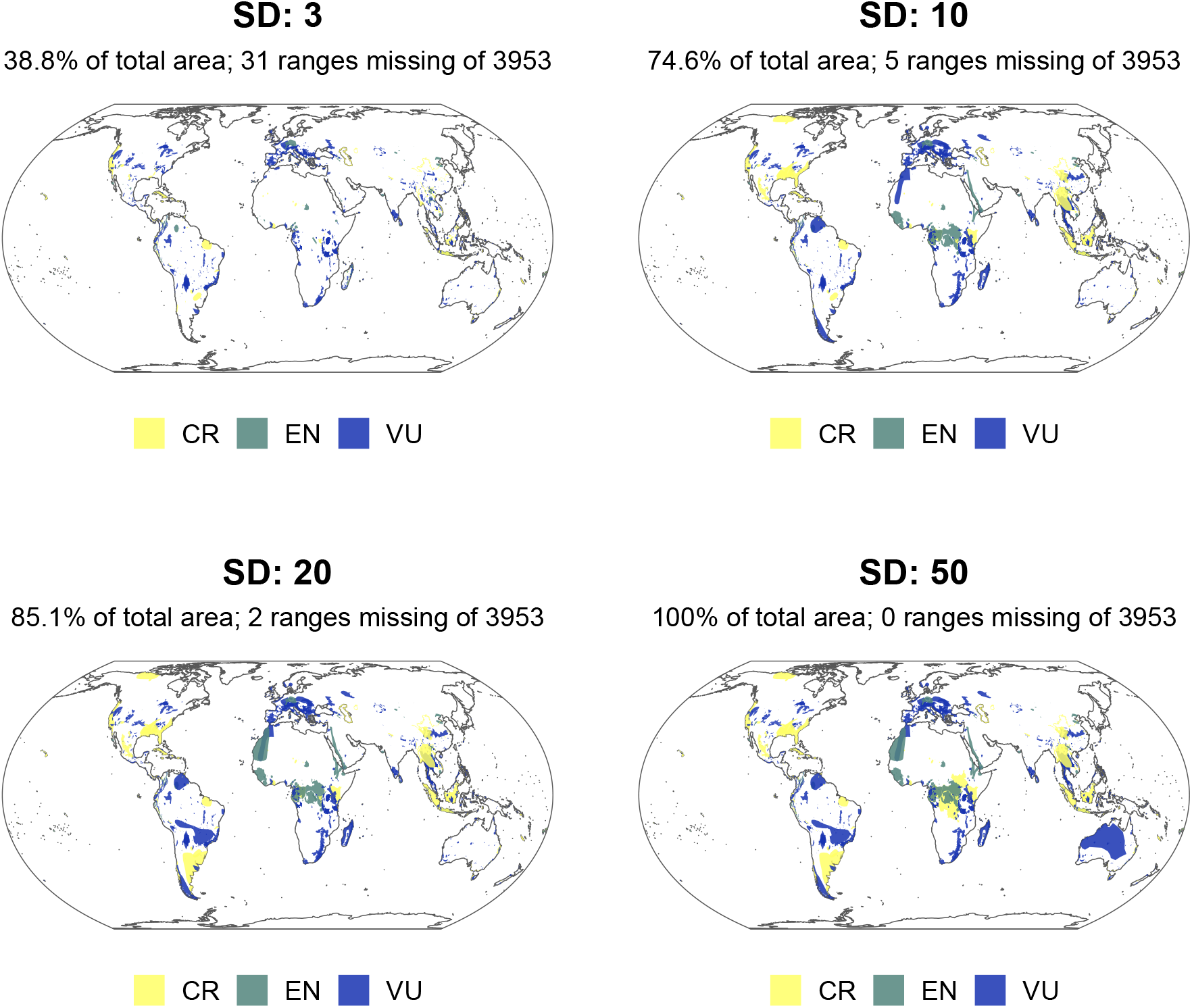
Sensitivity analysis of IUCN Red List ranges. Plots show result of removing areas with 3, 10, 20, 50 standard deviations (SDs) higher than the mean of the data. Each plot reports the SD, the number of ranges removed, and the percentage of the total area the data now represent.

#### Presence thresholds

Following Martin *et al*. ^15^ and Brauneder *et al*. ^16^, we use a high threshold for species distribution models of >90% as for the modelled cold-water coral datasets used in this analysis (Table 2; see also Figure 7). However, two datasets, tropical moist forest and mangroves, required an alternative approach as they represent high-resolution data (10 or 30m grid cells) derived from satellite imagery ^38,46^. The 10 or 30m pixels are classified as presence/absence. When averaged to 30 arcseconds resolution, these data then present an effective percentage cover in the cell. Figure 4 below shows the result of applying four different thresholds to these data: 0.25, 0.5, 0.75, and 0.9. As the aggregated data do not constitute a probability distribution, we selected ≥ 0.5 as the threshold. For mangroves, the area occupied by 1km cells over this threshold approximated the reported global extent of mangroves ^38^.

**Figure 4.**
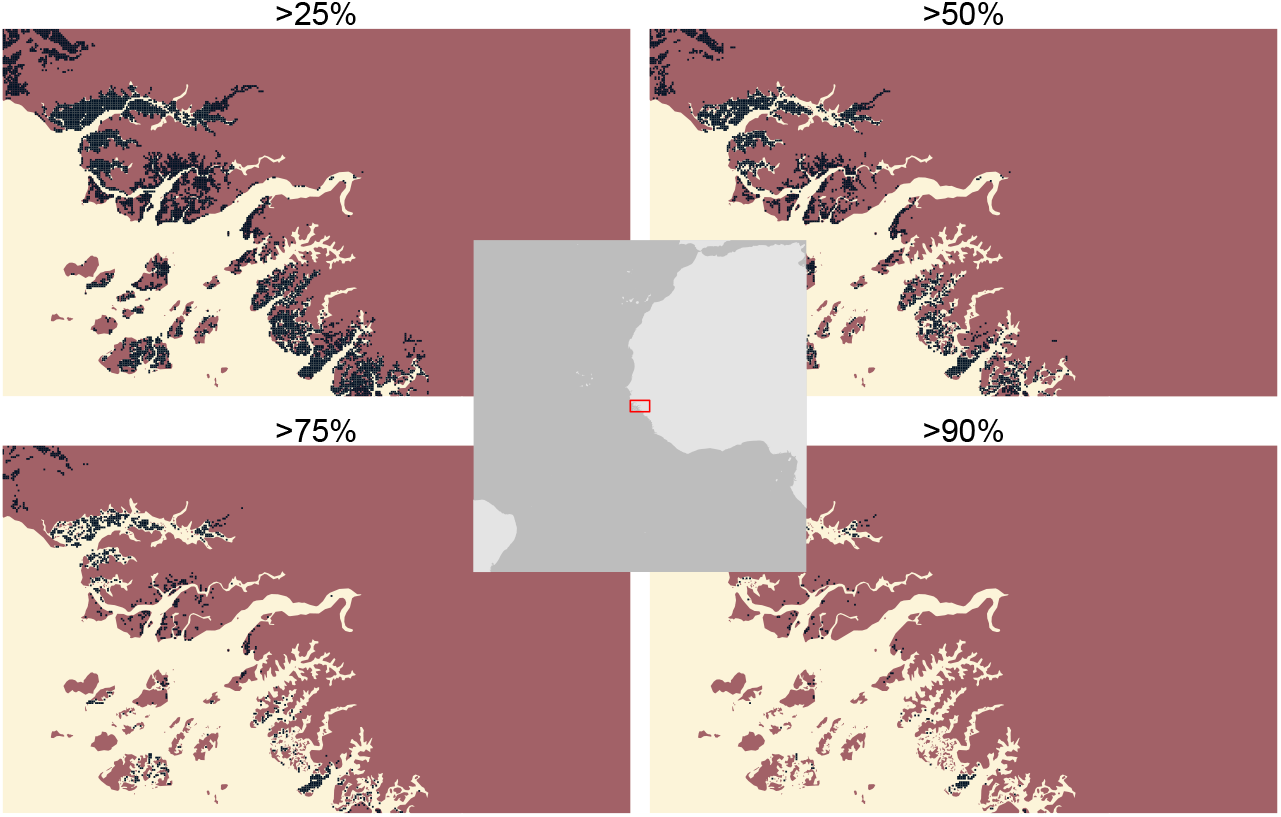
Sensitivity analysis of % presence cutoff values for mangrove data. Plots show effect of setting the % presence to >25,>50,>75,>90% on the output binary distribution. We select 50% for use in this analysis.

#### Feature coverage vs previous screening layer

38.15 million km^2^ has been added, 4.547 million km^2^ upgraded, 0.2528 million km^2^ downgraded, and 6.241 million km^2^ removed in the updated Critical Habitat screening layer. Table 8, Table 9, Table 10 and Table 11 detail the biodiversity features responsible for the changes. Figure 5 shows the composition of each biodiversity feature.

**Table 7.** States (including associated territories) with the largest areas of identified Critical Habitat.

| State | Likely CH | Potential CH | Total |
| --- | --- | --- | --- |
| Areas Beyond National Jurisdiction | 5,051 | 3,171 | 8,222 |
| United States of America (the) | 4,207 | 1,057 | 5,264 |
| Australia | 3,909 | 1,236 | 5,145 |
| Brazil | 3,976 | 478 | 4,454 |
| Russian Federation (the) | 4,067 | 186 | 4,253 |
| Canada | 3,972 | 78 | 4,050 |
| Indonesia | 1,568 | 816 | 2,383 |
| France | 1,616 | 679 | 2,295 |
| China | 1,659 | 127 | 1,785 |
| Congo (the Democratic Republic of the) | 1,473 | 77 | 1,551 |

*Note:*
Areas in km<sup>2</sup> x 10<sup>3</sup>.

**Table 8.** Biodiversity features contributing the five highest proportions to the area added as Potential or Likely Critical Habitat in the update.

| Biodiversity feature | Area added | % | Total feature area | % |
| --- | --- | --- | --- | --- |
| IMMAs under criteria C1, C2 and C3 | 11,367 | 29.80% | 12,812 | 88.72% |
| Intact Forest Landscapes | 7,900 | 20.71% | 11,908 | 66.34% |
| IMMAs under criterion A | 7,261 | 19.04% | 8,447 | 85.96% |
| IMMAs under criterion B2 | 4,679 | 12.27% | 5,462 | 85.66% |
| IMMAs under criterion D2 | 4,436 | 11.63% | 5,159 | 85.97% |

*Note:*
Areas in km<sup>2</sup> x 10<sup>3</sup>. IMMA: Important Marine Mammal Area. Percentages may add up to more than 100

**Table 9.** Biodiversity features contributing the five highest proportions to the area upgraded from Potential to Likely Critical Habitat in the update.

| Biodiversity feature | Area upgraded | % | Total feature area | % |
| --- | --- | --- | --- | --- |
| Tropical moist forest | 2,245 | 49.37% | 9,037 | 24.84% |
| Ever-wet tropical forests | 2,073 | 45.59% | 5,743 | 36.10% |
| IBAs under criterion A1 | 1,480 | 32.56% | 14,070 | 10.52% |
| Intact Forest Landscapes | 1,419 | 31.21% | 11,908 | 11.92% |
| IBAs under criterion A3 | 668.3 | 14.70% | 4,174 | 16.01% |

*Note:*
Areas in km<sup>2</sup> x 10<sup>3</sup>. IBA: Important Bird and Biodiversity Area. Percentages may add up to more than 100

**Table 10.** Five largest areas of biodiversity features downgraded from Likely to Potential Critical Habitat in the update. Names do not fully align with features in updated layer.

| Biodiversity feature | Area<br>downgraded | % | Previous<br>feature area | % |
| --- | --- | --- | --- | --- |
| KBA: 1 (CR/EN) | 3,237 | 27.56% | 69.68 | 2.15% |
| IBA: A3 | 5,048 | 25.39% | 64.21 | 1.27% |
| KBA: 2a | 1,547 | 21.76% | 55.02 | 3.56% |
| Mangroves | 487.2 | 13.45% | 34.01 | 6.98% |
| Warm-water coral reefs | 949.9 | 7.31% | 18.48 | 1.94% |

*Note:*
Areas in km<sup>2</sup> x 10<sup>3</sup>. CR: Critically Endangered; EN: Endangered; KBA: Key Biodiversity Area; IBA: Important Bird and Biodiversity Area. Percentages may add up to more than 100

**Table 11.** Five largest areas of biodiversity features removed as Critical Habitat in the update. Names do not fully align with features in updated layer.

| Biodiversity feature | Area<br>removed | % | Previous<br>feature area | % |
| --- | --- | --- | --- | --- |
| Tropical dry forest | 2,380 | 16.66% | 1,039 | 43.66% |
| PA (endemic/restricted range sp.) | 3,647 | 15.03% | 938.2 | 25.73% |
| KBA: 1 (CR/EN) | 3,237 | 12.92% | 806.3 | 24.91% |
| PA (CR/EN sp.) | 3,761 | 12.72% | 794.1 | 21.11% |
| Soft cold-water coral (Octocorals) | 1,226 | 7.60% | 474.5 | 38.72% |

*Note:*
Areas in km<sup>2</sup> x 10<sup>3</sup>. CR: Critically Endangered; EN: Endangered; PA: protected area; KBA: Key Biodiversity Area. Percentages may add up to more than 100

**Figure 5.**
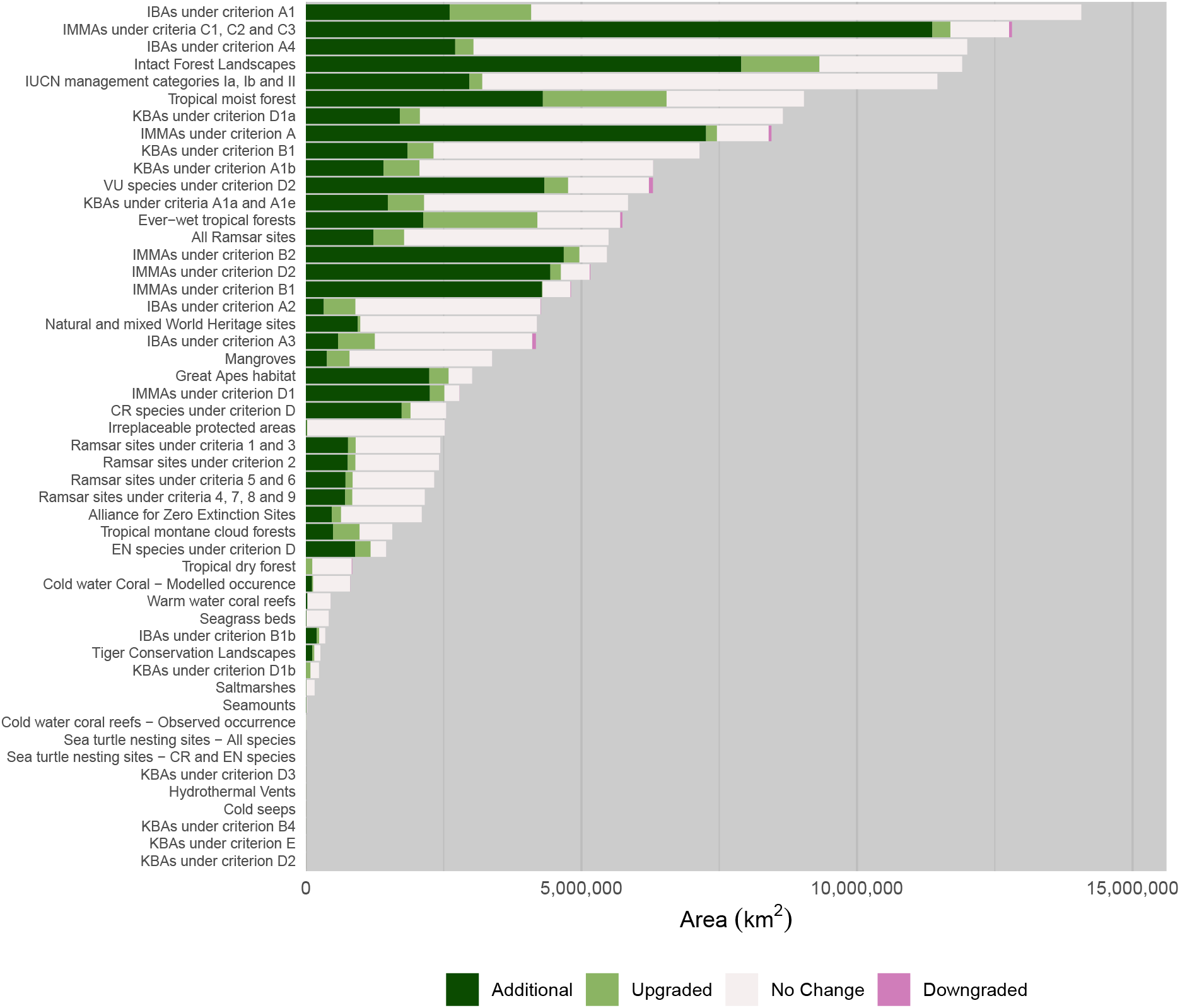
Critical Habitat (Likely and Potential) areas per biodiversity feature, split by whether the area is additional, upgraded, unchanged, or downgraded.

Important Marine Mammal Areas (IMMAs) dominate contributions to added Potential and Likely Critical Habitat (Table 8), adding 11.37 million km^2^ for IMMAs designated under criteria C1, C2 and C3 alone. Intact Forest Landscapes, an extensive biodiversity feature added in the update (see Supplementary Information for their inclusion justification), also contributes a significant area. For areas upgraded from Potential Critical Habitat to Likely Critical Habitat (Table 9), forest features dominate: over half of the area upgraded contain areas that are Likely Critical Habitat due to the presence of tropical moist forest. Just under half of all areas upgraded were previously Potential Critical Habitat due to the presence of tropical dry forest.

Areas downgraded or removed from the screening layer tend to be an order of magnitude lower than those upgraded or added (Table 10 and Table 11). Some of the features shown to have had area removed in the update are derived from the World Database on Protected Areas (WDPA) and World Database of Key Biodiversity Areas (WDKBA), two datasets that are regularly updated, with records both added and removed. Protected areas, for example, are often downgraded, downsized, or even degazetted ^50^ and these changes would be reflected in the screening layer. However, some features used the same data as previously, which warranted further inspection: 43.66% of tropical dry forest has been removed from the layer, alongside 38.72% of cold-water coral. Both appear to be small errors in the production of the previous screening layer. Figure 6 shows a comparison between the original input data, used in this layer and previously, alongside how the data were aggregated to a lower resolution in the previous layer and now. The method employed here appears to better reflect the higher resolution data but would reduce the overall feature area when compared to previously. For octocorals, the previous methodology considered values of **>90%** as presences for the species distribution models, but Figure 7 shows that it is more likely that the analysis classified cells ≥ 90% as presences. In this update, we have maintained a threshold of >90%, which has consequently removed some areas from the layer.

**Figure 6.**
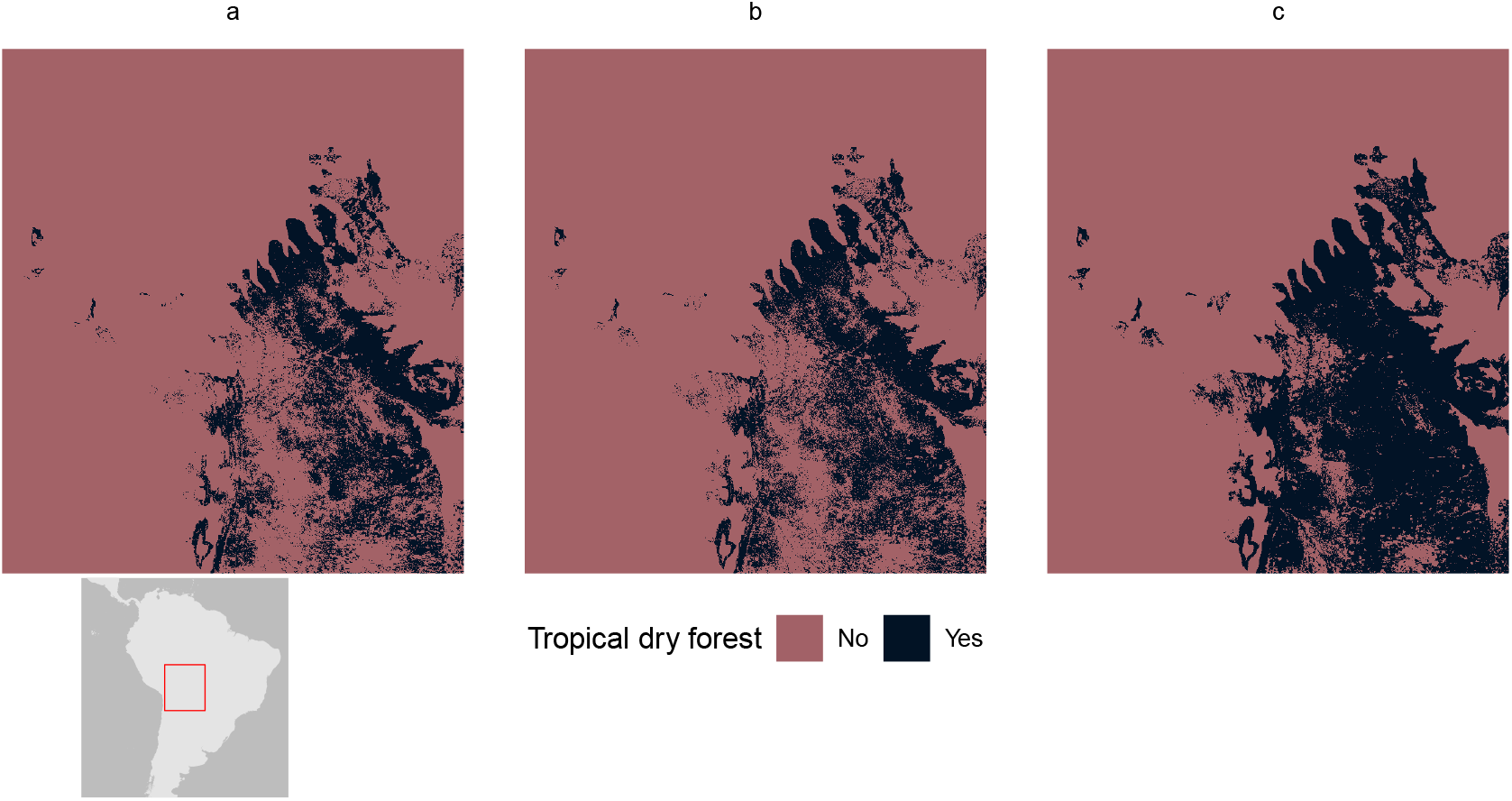
Differences in tropical dry forest coverage between (a) 2024 aggregation method, (b) original 500m resolution, and (c) 2018 aggregation method.

**Figure 7.**
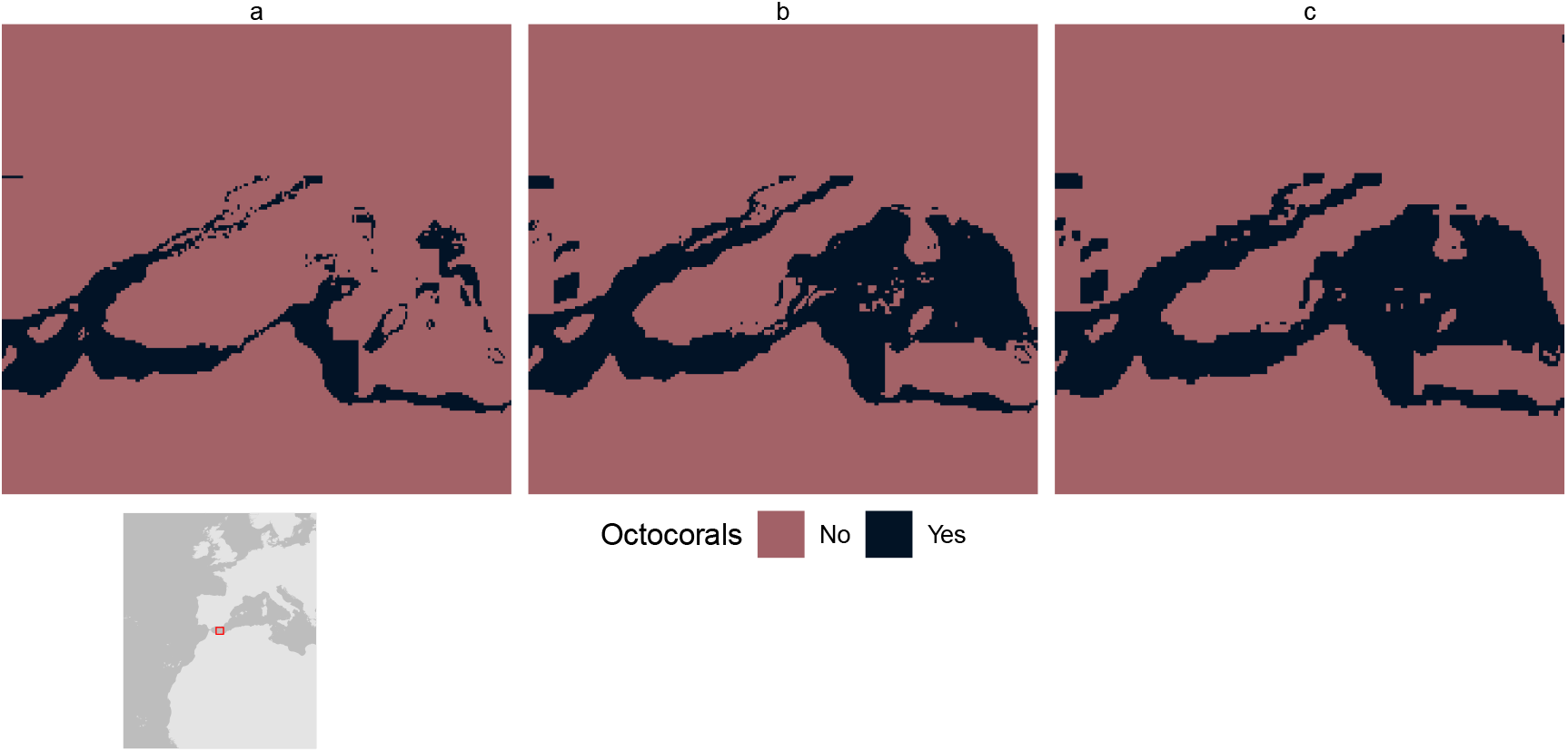
Differences in octocoral coverage between (a) 2024 threshold (>90%), (b) >=90% threshold, and (c) 2018 octocoral coverage.

### Usage notes

As noted by Martin *et al*. ^15^ and Brauneder *et al*. ^16^, the global Critical Habitat screening layer does not replace detailed on-the-ground Critical Habitat assessments. Reference to Critical Habitat here refers specifically to **areas in the screening layer identified as Likely or Potential Critical Habitat**, and does not indicate confirmed, on-the-ground, Critical Habitat. The data allow users to identify areas that need further investigation, including full Critical Habitat assessment, and to help direct impact mitigation efforts and conservation action.

We recommend working mostly with the raster data. While the polygon layer allows for more information to be stored about the relevant biodiversity triggers, it is susceptible to misuse if users mistake it for actual vector data and start performing advanced spatial operations on the layer.

Due to precision restrictions inherent with **R**, the number of biodiversity feature triggers that this workflow can handle is ∼66 (this analysis uses 54, see Table 2).

## Code availability

The code used to compile this update of the global Critical Habitat screening layer is publicly available through the *Zenodo* repository ^51^. The analysis is split across seven **R** scripts:

- *0 spatial_processing_functions.R*. Provides custom spatial processing functions required (see Methods).
- *1.1 Red_List_Preprocessing.R*. Initial processing of the IUCN Red List data, including filtering out species ranges with areas larger than a set amount of standard deviations above the mean (see Technical Validation).
- *1.2 Data_Preprocessing_Raster.R*. Preprocessing for the few datasets not available in vector format to produce standardised binary rasters for the next stage of the analysis.
- *1.3 Data_Preprocessing_Vector.R*. Preprocessing for vector input datasets to produce flat, dissolved layers for the next stage of the analysis.
- *2 Create_Drill_Down_Critical_Habitat_Raster_Layer.R*. Contains the lion’s share of processing. Converts the input flat vector layers to binary raster data and combines with the existing binary raster data in a process akin to the *Combine* tool in ArcGIS to produce a layer with a unique value for every resultant combination of input datasets. The script also produces the accompanying raster attribute table (RAT).
- *3 Create_Basic_Critical_Habitat_Raster_Layer.R*. Converts the detailed raster data into the basic Critical Habitat file.
- *4 Create_Drill_Down_Polygons.R*. Polygonises the drill down raster data to produce a vector layer that allows more information to be stored in the file than allowed in the spatial grid + RAT.

We regret that we cannot provide the full suite of input data due to copyright restrictions. All data not freely accessible through the UNEP-WCMC data portal are available on request from their respective owners.

## Footnotes

1 Reference to Critical Habitat in this manuscript refers specifically to *Likely or Potential* Critical Habitat. This **does not indicate** confirmed Critical Habitat, which requires ground-truthed assessment.

