## Supplementary Information for "An update to the global Critical Habitat screening layer"

#### ABSTRACT

The International Finance Corporation (IFC) defines Critical Habitat in Performance Standard 6 (PS6) as high biodiversity value areas requiring net biodiversity gain for projects. We present an updated global screening layer of Critical Habitat aligned with IFC's 2019 guidance. This layer derives from global datasets covering 53 biodiversity features, categorized as 'Likely' or 'Potential' Critical Habitat based on agreement with IFC criteria and data suitability. Analysis indicates 52.77 million km$^2$ (10.4%) and 15.94 million km$^2$ (3.2%) of the globe can be considered Likely and Potential Critical Habitat respectively, with the remaining 86.4% not overlapping with assessed biodiversity features. This represents a significant increase over previously identified 5% and 2% for Likely and Potential Critical Habitat. Likely Critical Habitat was dominated by Important Bird and Biodiversity Areas, Intact Forest Landscapes, and protected areas; Potential Critical Habitat by Important Marine Mammal Areas and ranges of IUCN Vulnerable species. Our results can help businesses prioritize impact avoidance and identify opportunities by screening potential development sites for biodiversity features.

#### 1 Spherical geometries with the S2 library

To maintain the highest standards of accuracy, spatial processing is carried out with the `sf` package in **R**, using the S2 spherical geometry library. This allows spatial processing, e.g. intersections and buffers, using Great Circle distances.

However, many conservation datasets are set up for planar geometry and are not valid on the sphere. To account for this, data are forced to be valid on the sphere through two methods. The first is light touch, calling the function `st_make_valid` from the `sf` package to remove duplicate vertices etc. The second, more brute force approach, is to transform the data to projected coordinates, e.g. Mollweide equal-area, and call `st_simplify` with rising imprecision until the geometry is valid on the sphere. Generally, where the latter approach was required, geometries were made valid before the precision rose above 1-km resolution (`dTolerance = 1000`).

#### 2 Unioning geometries

In order to compute accurate area measurements for polygon input features and remove duplicate records from point data, all data are unioned using `st_union` from the `sf` package. However, using `st_union` for spherical spatial processing becomes extremely inefficient (e.g. days to compute) with large, high precision datasets. For this reason, an arbitrary complexity limit was used, above which planar intersection with `unary_union` from the `geos` package was performed. This limit was set at 10,000,000 vertices: only a handful of datasets are more detailed than this, and it speeds up processing markedly with limited impact on accuracy.

#### 3 Cold seeps

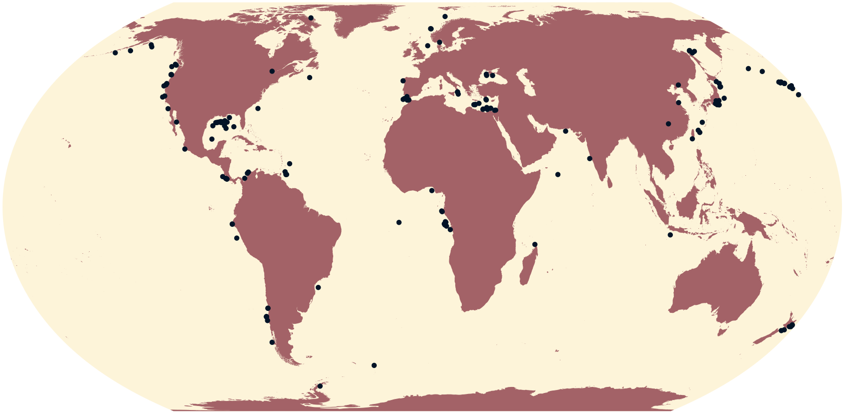

**Figure 1.** Global distribution of cold seeps.

- **Data type**: Point
- **Number of records**: 188
- **Original CRS**: WGS 84
- **Critical Habitat type**: Likely/Potential
- **IFC PS6 Criteria triggered**: C4 (Highly Threatened or Unique Ecosystems) and C5 (Key Evolutionary Processes)/C2 (Endemic and Restricted-range Species)
- **Source**: Ramirez-Llodra 2020[1]

***Justification and alignment with IFC PS6 criteria***
See Martin *et al.* 2015[2]. No change to data.

***How the data are filtered***
No filtering of data from source.

***How the data are processed***
Data are unioned to remove duplicates (see Section 2).

### 4 Cold water corals

#### 4.1 Modelled occurrence

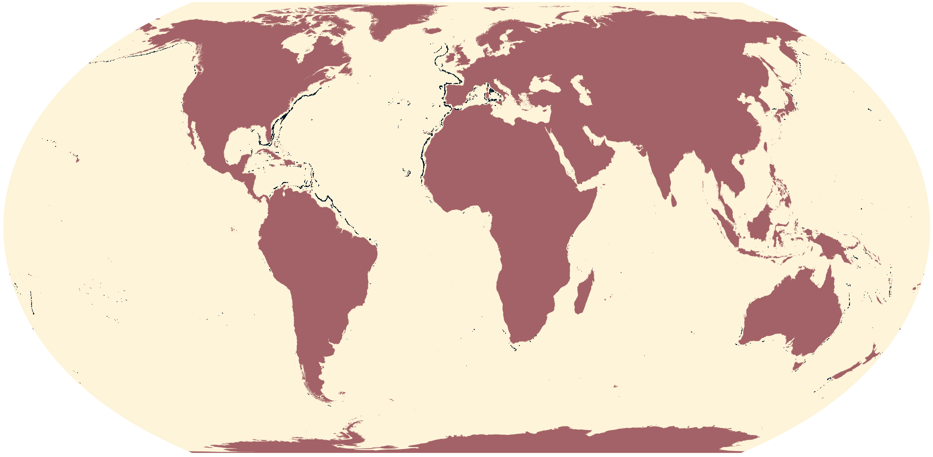

**Figure 2.** Global modelled distribution of cold water corals.

- **Data type**: Raster
- **Original CRS**: WGS_1984_Cylindrical_Equal_Area
- **Original resolution**: 1,000
- **Extent**: -20,040,000-6,364,00020,040,0006,364,000
- **Gridded area of feature at 1-km resolution**: 802,500
- **Critical Habitat type**: Potential
- **IFC PS6 Criteria triggered**:
  - C4 (Highly Threatened or Unique Ecosystems)
  - C5 (Key Evolutionary Processes)
- **Source**: Yesson *et al.* 2012[3]

***Justification and alignment with IFC PS6 criteria***
See Martin *et al.* 2015[2]. No change to data.

***How the data are processed***
- As per Martin *et al.* 2015[2], raster reclassified so that `>90 = 1` and `<=90 = 0`.
- Raster projected to WGS 84 using nearest neighbour resampling.

#### 4.2 Observed occurence

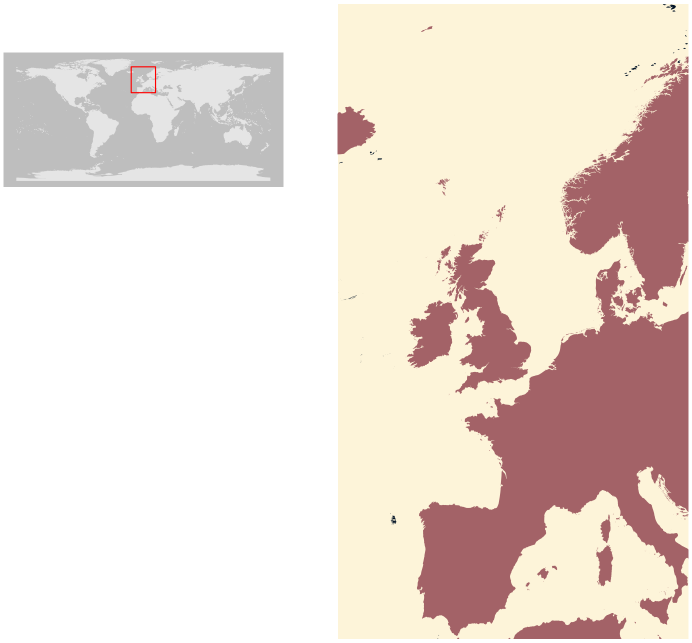

**Figure 3.** Global observed distribution of cold water corals - polygons (inset: North Atlantic and North Sea).

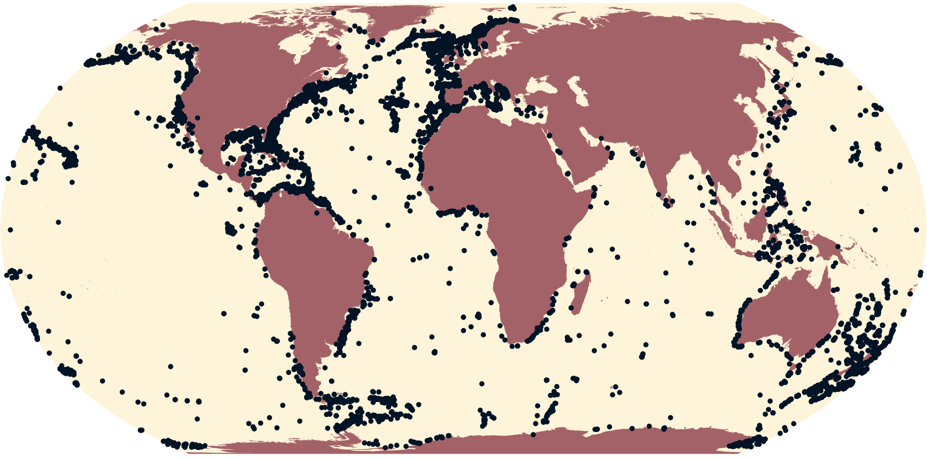

**Figure 4.** Global observed distribution of cold water corals - points.

- **Data type**: Point and polygon
- **Number of records**: 15,430 (point) and 37 (polygon)
- **Original CRS**: WGS 84
- **Area (polygons)**: 3,932 $km^2$
- **Critical Habitat type**: Likely
- **IFC PS6 Criteria triggered**:

  - C4 (Highly Threatened or Unique Ecosystems)
  - C5 (Key Evolutionary Processes)
- **Source**: Freiwald *et al.* 2021[4]

***Justification and alignment with IFC PS6 criteria***

See Martin *et al.* 2015[2]. No change to data.

***How the data are filtered***

No filtering of data from source.

***How the data are processed***

- Invalid S2 polygon geometries (see Section 1) are made valid.
- Data are unioned to remove overlap (polygons) and duplicates (points) (see Section 2).

#### 5 Ever-wet tropical forest

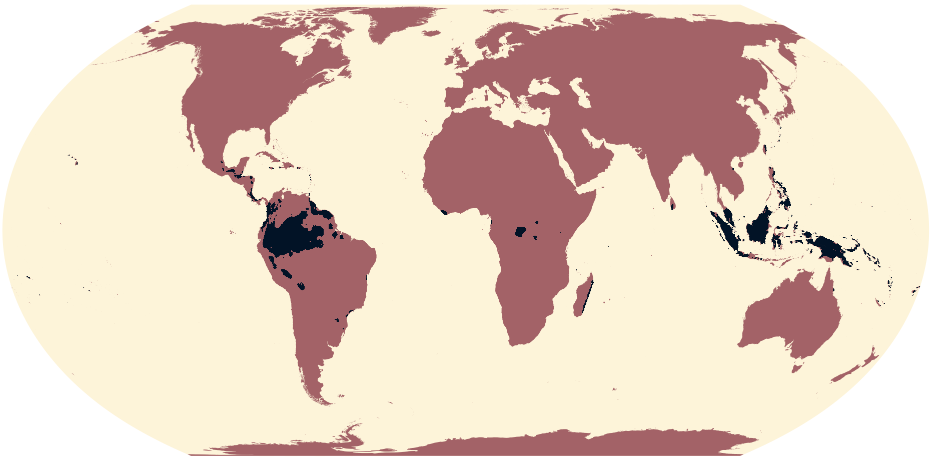

**Figure 5.** Global distribution of ever-wet tropical forests.

- **Data type**: Raster
- **Original CRS**: CYLINDRICAL
- **Original resolution**: 3,911
- **Extent**: -20,040,000-3,559,00020,040,0003,257,000
- **Gridded area of feature at 1-km resolution**: 5,743,000
- **Critical Habitat type**: Potential
- **IFC PS6 Criteria triggered**: C4 (Highly Threatened or Unique Ecosystems)
- **Source**: Underwood *et al* 2014[5]

***Justification and alignment with IFC PS6 criteria***
See Brauneder *et al.* 2018[6]. No change to data.

***How the data are processed***
- Reclassify Current ever-wet zones to 1, all other categories to 0.
- Simple disaggregation to desired resolution, i.e. each cell split into several cells, all of which are given the same value as the parent cell.

### 6 Hydrothermal vents

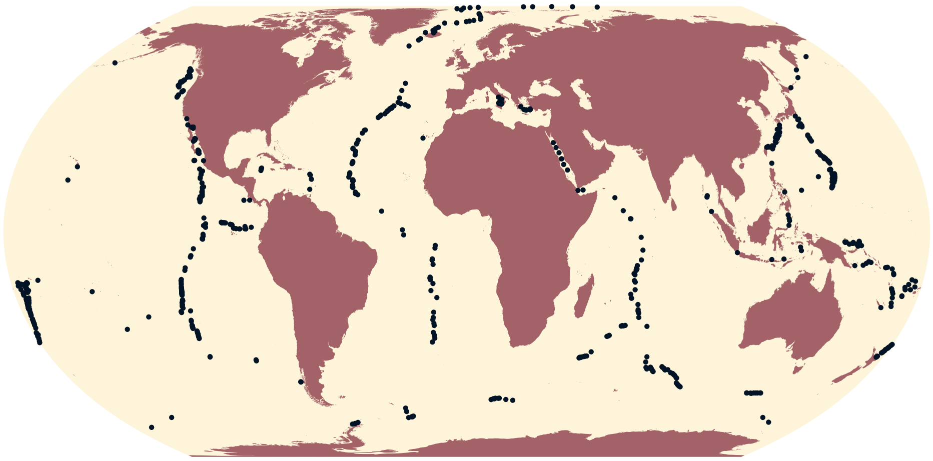

**Figure 6.** Global distribution of active hydrothermal vents.

- **Data type**: Point
- **Number of records**: 721; of which 666 are active
- **Original CRS**: WGS 84
- **Critical Habitat type**: Likely (if active) / Potential
- **IFC PS6 Criteria triggered**: C2 (Endemic and Restricted-range Species) and C5 (Key Evolutionary Processes) (if active) / C4 (Highly Threatened or Unique Ecosystems)
- **Source**: Beaulieu and Szafrański 2020[7]

***Justification and alignment with IFC PS6 criteria***
See Martin *et al.* 2015[2]. No change to data.

***How the data are filtered***
Records whose `Activity` is classified as `active,inferred` or `active, confirmed`.

***How the data are processed***
Data are unioned to remove duplicates (see Section 2).

### 7 Key Biodiversity Areas (KBAs)

- **Data type**: Point and polygon
- **Number of records**:317 (point) and 16,230 (polygon)
- **Original CRS**: WGS 84
- **Area (polygons - reported)**: 22,130,000 km$^2$
- **Area (point - reported)**: 0 km$^2$
- **Source**: BirdLife International 2024[8]

#### 7.1 Alliance for Zero Extinction Sites (AZEs)

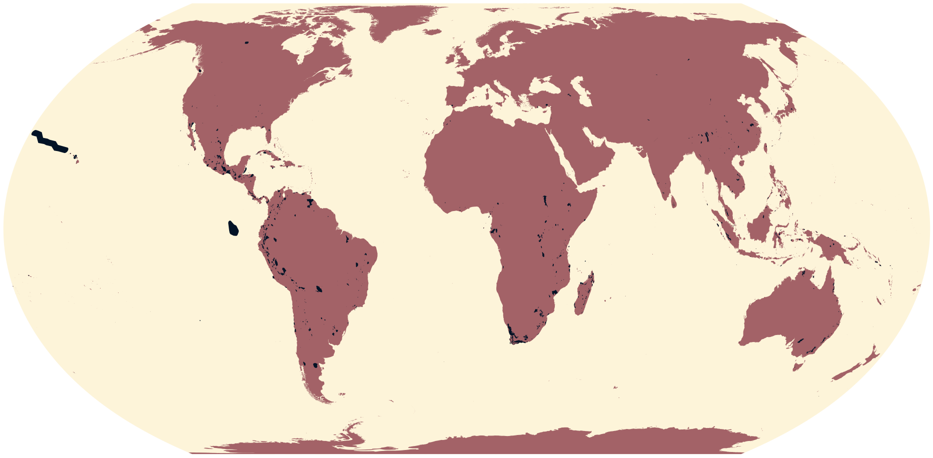

**Figure 7.** Global distribution of Alliance for Zero Extinction sites.

- **Number of records**: 3 (point) and 970 (polygon)
- **Area (including buffered point data)**: 1,804,000 km$^2$
- **Critical Habitat type**: Likely
- **IFC PS6 Criteria triggered**:
  - C1 (Critically Endangered and Endangered Species)
  - C2 (Endemic and Restricted-range Species)
  - C3 (Migratory and Congregatory Species)

***Justification and alignment with IFC PS6 criteria***

See Martin *et al.* 2015[2] and Brauneder *et al.* 2018[6]. Updated data.

***How the data are filtered***

- Sites with `AzeStatus = confirmed`.
- All polygons, and point data with reported areas.

***How the data are processed***

- Invalid S2 geometries (see Section 1) are made valid.
- Data are unioned to remove overlap (see Section 2).

##### 7.2 KBAs under criteria A1a and A1e

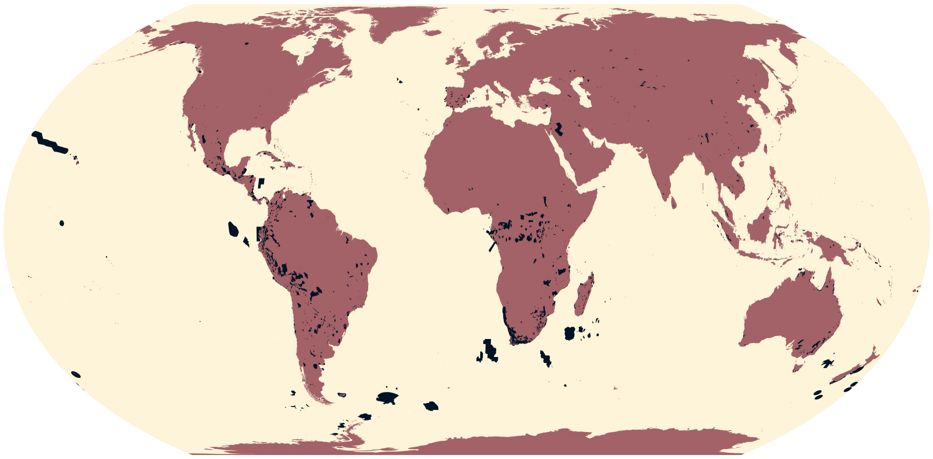

**Figure 8.** Global distribution of Key Biodiversity Areas under criteria A1a and A1e.

- **Number of records**: 6 (point) and 2,205 (polygon)
- **Area (including buffered point data)**: 5,611,000 km$^2$
- **Critical Habitat type**: Likely
- **IFC PS6 Criteria triggered**: C1 (Critically Endangered and Endangered Species)

***Justification and alignment with IFC PS6 criteria***

Much of the rationale for the updated 2019 IFC Guidance Note 6 was to align Critical Habitat criteria thresholds with the since published *Global Standard for the Identification of Key Biodiversity Areas*[9,10,11]. Building on the justifications set out in Martin *et al.* 2015[2] and Brauneder *et al.* 2018[6], KBAs designated under KBA Criteria A1a ($\geq$ 0.5% of global population size and $\geq$ 5 reproductive units (RU) of a CR/EN species) and KBA Criteria A1e (Effectively the entire population size of a CR/EN species) align directly with thresholds GN72(a) and GN72(c) of GN2019 GN72[9].

***How the data are filtered***

- Sites with `KbaStatus = confirmed`.
- Sites designated under:
  - KBA Criteria A1a ($\geq$ 0.5% of global population size and $\geq$ 5 reproductive units (RU) of a CR/EN species); and
  - KBA Criteria A1e (Effectively the entire population size of a CR/EN species).
- All polygons, and point data with reported areas.

***How the data are processed***

- Invalid S2 geometries (see Section 1) are made valid.
- Point data buffered to reported area, `sitarea`.
- Data are unioned to remove overlap (see Section 2).

##### 7.3 KBAs under criterion A1b

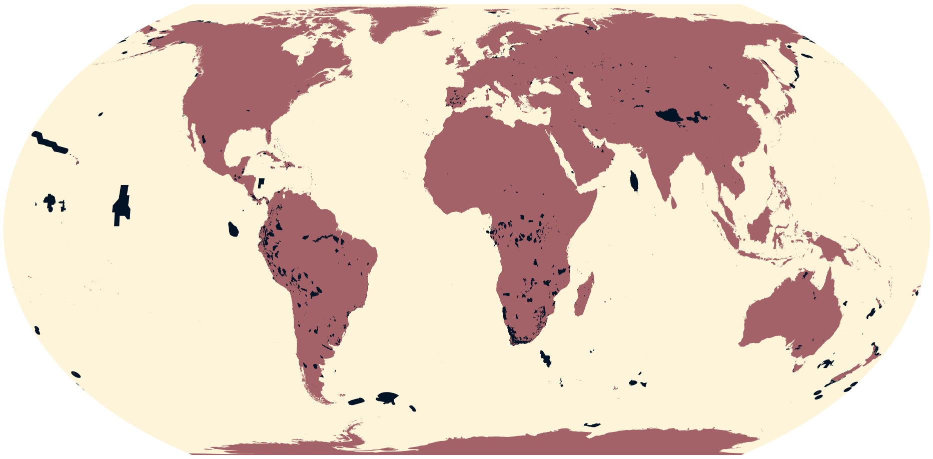

**Figure 9.** Global distribution of Key Biodiversity Areas under criterion A1b.

- **Number of records**: 14 (point) and 1,454 (polygon)
- **Area (including buffered point data)**: 6,125,000 $km^2$
- **Critical Habitat type**: Potential
- **IFC PS6 Criteria triggered**: C1 (Critically Endangered and Endangered Species)

***Justification and alignment with IFC PS6 criteria***

Much of the rationale for the updated 2019 IFC Guidance Note 6 was to align Critical Habitat criteria thresholds with the since published *Global Standard for the Identification of Key Biodiversity Areas*[9,10,11]. Building on the justifications set out in Martin *et al.* 2015[2] and Brauneder *et al.* 2018[6], KBAs designated under KBA Criterion A1b ($\geq$ 1.0% of global population size and $\geq$ 10 RU of a VU species) align with threshold GN72(b) of GN2019[9] by representing globally important concentrations of IUCN Red-listed VU species.

***How the data are filtered***

- Sites with `KbaStatus = confirmed`.
- Sites designated under KBA Criterion A1b ($\geq$ 1.0% of global population size and $\geq$ 10 RU of a VU species).
- All polygons, and point data with reported areas.

***How the data are processed***

- Invalid S2 geometries (see Section 1) are made valid.
- Point data buffered to reported area, `sitarea`.
- Data are unioned to remove overlap (see Section 2).

##### 7.4 KBAs under criterion A2a

No data present.

##### 7.5 KBAs under criterion A2b

No data present.

##### 7.6 KBAs under criterion B1

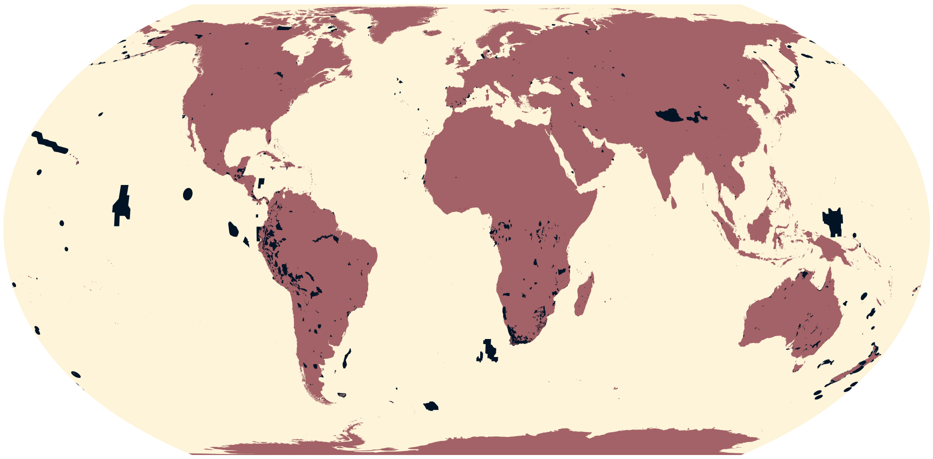

**Figure 10.** Global distribution of Key Biodiversity Areas under criterion B1.

- **Number of records**: 2 (point) and 1,622 (polygon)
- **Area (including buffered point data)**: 6,943,000 km$^2$
- **Critical Habitat type**: Likely
- **IFC PS6 Criteria triggered**: C2 (Endemic and Restricted-range Species)

***Justification and alignment with IFC PS6 criteria***

Much of the rationale for the updated 2019 IFC Guidance Note 6 was to align Critical Habitat criteria thresholds with the since published *Global Standard for the Identification of Key Biodiversity Areas*[9,10,11]. Building on the justifications set out in Martin *et al.* 2015[2] and Brauneder *et al.* 2018[6], KBAs designated under KBA Criterion B1 (Site regularly holds $\geq$ 10% of the global population size AND $\geq$ 10 reproductive units of a species) align directly with threshold GN75(a) of GN2019[9].

***How the data are filtered***

- Sites with `KbaStatus = confirmed`.
- Sites designated under KBA Criterion B1 (Site regularly holds $\geq$ 10% of the global population size AND $\geq$ 10 reproductive units of a species).
- All polygons, and point data with reported areas.

***How the data are processed***

- Invalid S2 geometries (see Section 1) are made valid.
- Point data buffered to reported area, `sitarea`.
- Data are unioned to remove overlap (see Section 2).

##### 7.7 KBAs under criterion B4

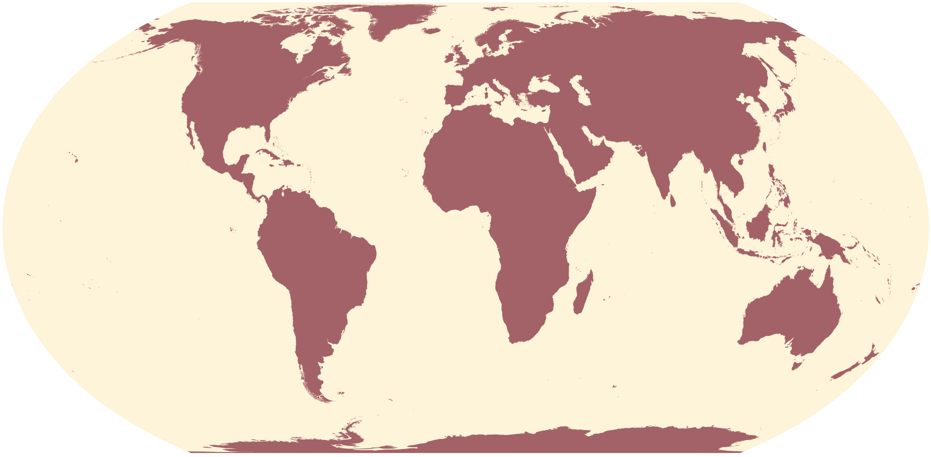

**Figure 11.** Global distribution of Key Biodiversity Areas under criterion B4.

- **Number of records**: 0 (point) and 2 (polygon)
- **Area (including buffered point data)**: 218.6 $km^2$
- **Critical Habitat type**: Potential
- **IFC PS6 Criteria triggered**: C4 (Highly Threatened or Unique Ecosystems)

***Justification and alignment with IFC PS6 criteria***

Much of the rationale for the updated 2019 IFC Guidance Note 6 was to align Critical Habitat criteria thresholds with the since published *Global Standard for the Identification of Key Biodiversity Areas*[9,10,11]. Building on the justifications set out in Martin *et al.* 2015[2] and Brauneder *et al.* 2018[6], KBAs designated under KBA Criterion B4 (Site holds $\geq$ 20% of the global extent of an ecosystem type) align with threshold GN80(a) of GN2019[9]. The IUCN Red List of Ecosystems does not yet have full global coverage, but KBA Criterion B4 aligns by providing a higher threshold ($\geq$ 20%) than GN80(a) ($\geq$ 5%) to account for lack of threat classifications.

***How the data are filtered***

- Sites with `KbaStatus = confirmed`.
- Sites designated under KBA Criterion B4 (Site holds $\geq$ 20% of the global extent of an ecosystem type).
- All polygons, and point data with reported areas.

***How the data are processed***

- Invalid S2 geometries (see Section 1) are made valid.
- Point data buffered to reported area, `sitarea`.
- Data are unioned to remove overlap (see Section 2).

##### 7.8 KBAs under criterion C

No data present.

##### 7.9 KBAs under criterion D1a

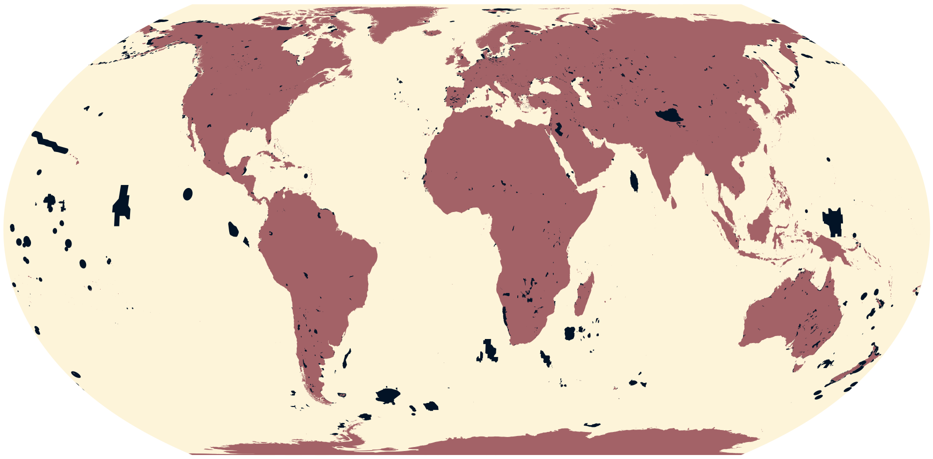

**Figure 12.** Global distribution of Key Biodiversity Areas under criterion D1a.

- **Number of records**: 24 (point) and 2,749 (polygon)
- **Area (including buffered point data)**: 8,434,000 km$^2$
- **Critical Habitat type**: Likely
- **IFC PS6 Criteria triggered**: C3 (Migratory and Congregatory Species)

***Justification and alignment with IFC PS6 criteria***

Much of the rationale for the updated 2019 IFC Guidance Note 6 was to align Critical Habitat criteria thresholds with the since published *Global Standard for the Identification of Key Biodiversity Areas*[9,10,11]. Building on the justifications set out in Martin *et al.* 2015[2] and Brauneder *et al.* 2018[6], KBAs designated under KBA Criterion D1a (Site holds an aggregation representing ≥ 1% of the global population size of a species, over a season, and during one or more key stages of its life cycle) align directly with threshold GN78(a) of GN2019[9].

***How the data are filtered***

- Sites with `KbaStatus = confirmed`.
- Sites designated under KBA Criterion D1a (Site holds an aggregation representing ≥ 1% of the global population size of a species, over a season, and during one or more key stages of its life cycle).
- All polygons, and point data with reported areas.

***How the data are processed***

- Invalid S2 geometries (see Section 1) are made valid.
- Point data buffered to reported area, `sitarea`.
- Data are unioned to remove overlap (see Section 2).

#### 7.10 KBAs under criterion D1b

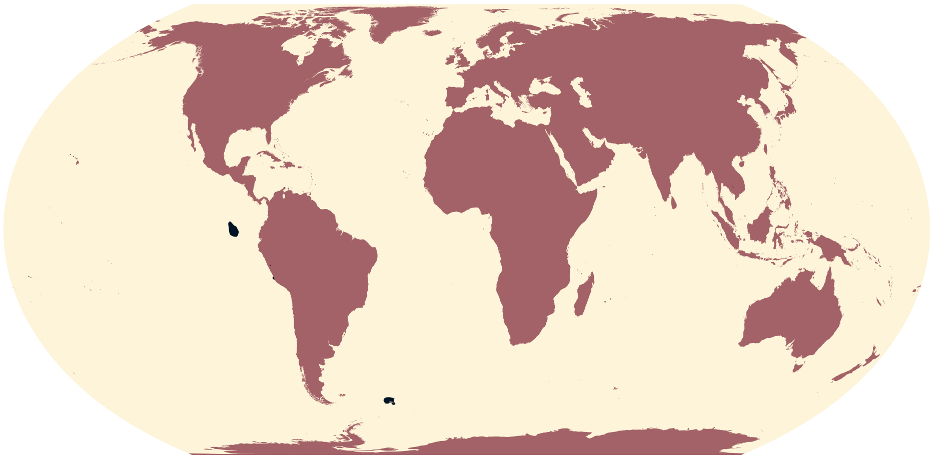

**Figure 13.** Global distribution of Key Biodiversity Areas under criterion D1b.

- **Number of records**: 0 (point) and 14 (polygon)
- **Area (including buffered point data)**: 235,300 km$^2$
- **Critical Habitat type**: Potential
- **IFC PS6 Criteria triggered**: C3 (Migratory and Congregatory Species)

***Justification and alignment with IFC PS6 criteria***

Much of the rationale for the updated 2019 IFC Guidance Note 6 was to align Critical Habitat criteria thresholds with the since published *Global Standard for the Identification of Key Biodiversity Areas*[9,10,11]. Building on the justifications set out in Martin *et al.* 2015[2] and Brauneder *et al.* 2018[6], KBAs designated under KBA Criterion D1b (Site holds a number of mature individuals that ranks the site among the largest 10 aggregations known for the species) align with threshold GN78(a) and GN78(b) of GN2019[9].

***How the data are filtered***

- Sites with `KbaStatus = confirmed`.
- Sites designated under KBA Criterion D1b (Site holds a number of mature individuals that ranks the site among the largest 10 aggregations known for the species).
- All polygons, and point data with reported areas.

***How the data are processed***

- Invalid S2 geometries (see Section 1) are made valid.
- Point data buffered to reported area, `sitarea`.
- Data are unioned to remove overlap (see Section 2).

#### 7.11 KBAs under criterion D2

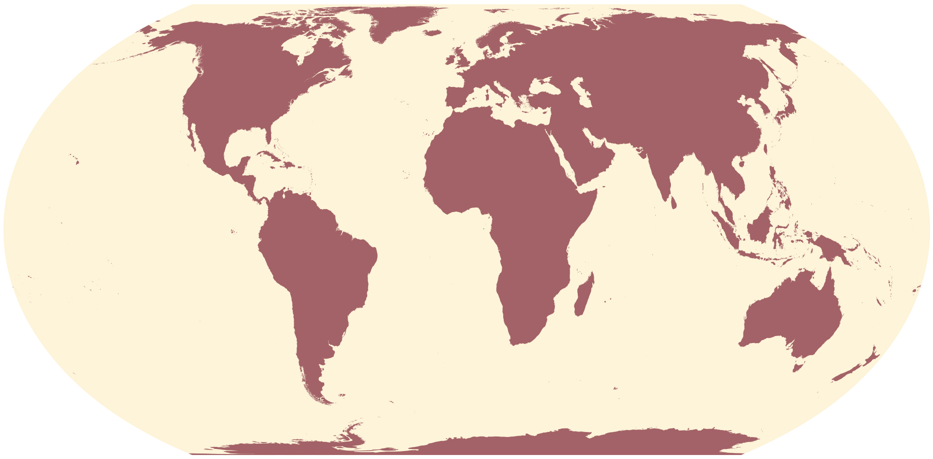

**Figure 14.** Global distribution of Key Biodiversity Areas under criterion D2.

- **Number of records**: 0 (point) and 1 (polygon)
- **Area (including buffered point data)**: 1.616 km$^2$
- **Critical Habitat type**: Likely
- **IFC PS6 Criteria triggered**: C3 (Migratory and Congregatory Species)

***Justification and alignment with IFC PS6 criteria***

Much of the rationale for the updated 2019 IFC Guidance Note 6 was to align Critical Habitat criteria thresholds with the since published *Global Standard for the Identification of Key Biodiversity Areas*[9,10,11]. Building on the justifications set out in Martin *et al.* 2015[2] and Brauneder *et al.* 2018[6], KBAs designated under KBA Criterion D2 (Site supports $\geq$ 10% of the global population size of one or more species during periods of environmental stress, for which historical evidence shows that it has served as a refugium in the past and for which there is evidence to suggest it would continue to do so in the foreseeable future) align directly with threshold GN78(b) of GN2019[9].

***How the data are filtered***

- Sites with `KbaStatus = confirmed`.
- Sites designated under KBA Criterion D2 (Site supports $\geq$ 10% of the global population size of one or more species during periods of environmental stress, for which historical evidence shows that it has served as a refugium in the past and for which there is evidence to suggest it would continue to do so in the foreseeable future).
- All polygons, and point data with reported areas.

***How the data are processed***

- Invalid S2 geometries (see Section 1) are made valid.
- Point data buffered to reported area, `sitarea`.
- Data are unioned to remove overlap (see Section 2).

##### 7.12 KBAs under criterion D3

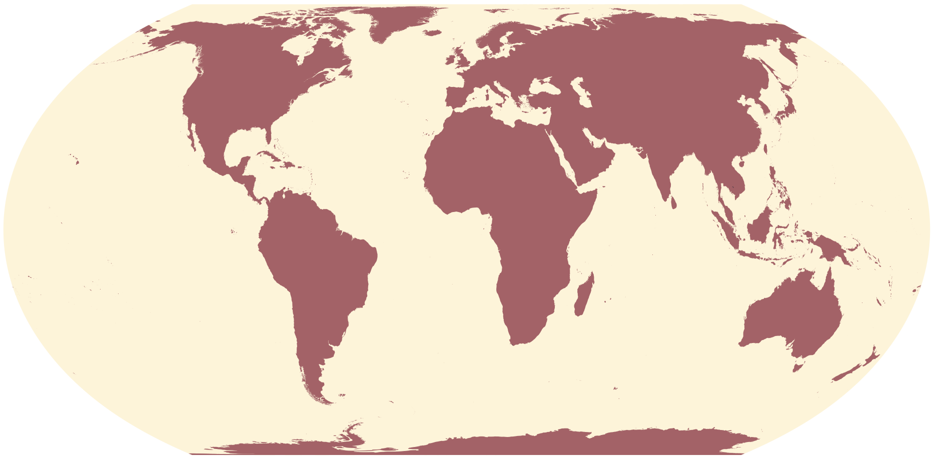

**Figure 15.** Global distribution of Key Biodiversity Areas under criterion D3.

- **Number of records**: 0 (point) and 3 (polygon)
- **Area (including buffered point data)**: 1.616 km$^2$
- **Critical Habitat type**: Potential
- **IFC PS6 Criteria triggered**: C3 (Migratory and Congregatory Species)

***Justification and alignment with IFC PS6 criteria***
Much of the rationale for the updated 2019 IFC Guidance Note 6 was to align Critical Habitat criteria thresholds with the since published *Global Standard for the Identification of Key Biodiversity Areas*[9,10,11]. Building on the justifications set out in Martin *et al.* 2015[2] and Brauneder *et al.* 2018[6], KBAs designated under KBA Criterion D3 (Site predictably produces propagules, larvae, or juveniles that maintain $\geq$ 10% of the global population size of a species) align with threshold GN78(a) and GN78(b) of GN2019[9].

***How the data are filtered***
- Sites with `KbaStatus = confirmed`.
- Sites designated under KBA Criterion D3 (Site predictably produces propagules, larvae, or juveniles that maintain $\geq$ 10% of the global population size of a species).
- All polygons, and point data with reported areas.

***How the data are processed***
- Invalid S2 geometries (see Section 1) are made valid.
- Point data buffered to reported area, `sitarea`.
- Data are unioned to remove overlap (see Section 2).

#### 7.13 KBAs under criterion E

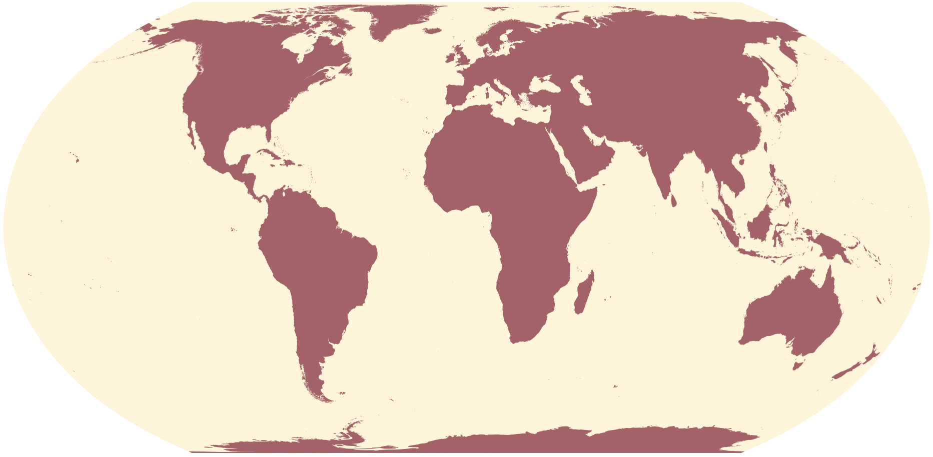

**Figure 16.** Global distribution of Key Biodiversity Areas under criterion E.

- **Number of records**: 0 (point) and 3 (polygon)
- **Area (including buffered point data)**: 26.92 $km^2$
- **Critical Habitat type**: Potential
- **IFC PS6 Criteria triggered**: C1 (Critically Endangered and Endangered Species)

***Justification and alignment with IFC PS6 criteria***

Much of the rationale for the updated 2019 IFC Guidance Note 6 was to align Critical Habitat criteria thresholds with the since published *Global Standard for the Identification of Key Biodiversity Areas*[9,10,11]. Building on the justifications set out in Martin *et al.* 2015[2] and Brauneder *et al.* 2018[6], KBAs designated under KBA Criterion E (Site has a level of irreplaceability of $\geq$ 0.90 (on a 0–1 scale), measured by quantitative spatial analysis, and is characterised by the regular presence of species with $\geq$ 10 reproductive units known to occur (or $\geq$ 5 units for EN or CR species)) align with threshold GN72(a) and GN72(c) of GN2019[9].

***How the data are filtered***

- Sites with `KbaStatus = confirmed`.
- Sites designated under KBA Criterion E (Site has a level of irreplaceability of $\geq$ 0.90 (on a 0–1 scale), measured by quantitative spatial analysis, and is characterised by the regular presence of species with $\geq$ 10 reproductive units known to occur (or $\geq$ 5 units for EN or CR species)).
- All polygons, and point data with reported areas.

***How the data are processed***

- Invalid S2 geometries (see Section 1) are made valid.
- Point data buffered to reported area, `sitarea`.
- Data are unioned to remove overlap (see Section 2).

### 8 Important Bird and Biodiversity Areas (IBAs)

- **Data type**: Point and polygon
- **Number of records**: 149 (point) and 13,720 (polygon)
- **Original CRS**: WGS 84
- **Area (polygons - reported)**: 19,180,000 km$^2$
- **Area (point - reported)**: 0 km$^2$
- **Source**: BirdLife International 2023[12]

#### 8.1 IBAs under criterion A1

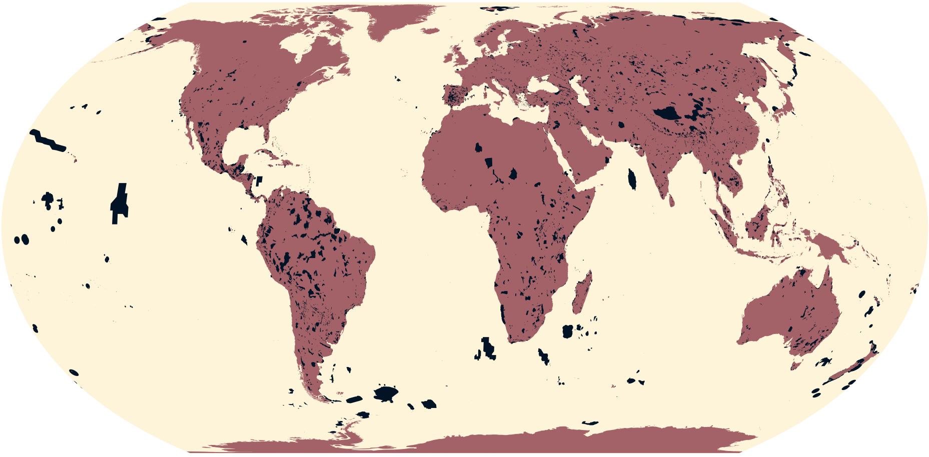

**Figure 17.** Global distribution of Important Bird and Biodiversity Areas under criterion A1.

- **Number of records**: 85 (point) and 7,711 (polygon)
- **Area (including buffered point data)**: 13,460,000 km$^2$
- **Critical Habitat type**: Likely
- **IFC PS6 Criteria triggered**: C1 (Critically Endangered and Endangered Species)

***Justification and alignment with IFC PS6 criteria***
See Martin *et al.* 2015[2] and Brauneder *et al.* 2018[6]. Updated data.

***How the data are filtered***
- Sites designated under IBA Criterion A1 (The site is known or thought regularly to hold significant numbers of a globally threatened species).
- All polygons, and point data with reported areas.

***How the data are processed***
- Invalid S2 geometries (see Section 1) are made valid.
- Point data buffered to reported area, `SitArea`.
- Data are unioned to remove overlap (see Section 2).

#### 8.2 IBAs under criterion A2

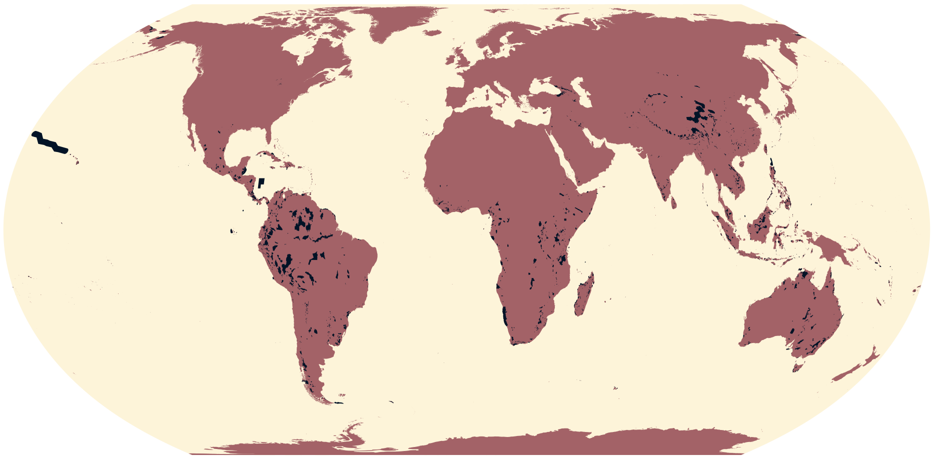

**Figure 18.** Global distribution of Important Bird and Biodiversity Areas under criterion A2.

- **Number of records**: 44 (point) and 2,754 (polygon)
- **Area (including buffered point data)**: 4,025,000 $km^2$
- **Critical Habitat type**: Potential
- **IFC PS6 Criteria triggered**: C2 (Endemic and Restricted-range Species)

***Justification and alignment with IFC PS6 criteria***

See Martin *et al.* 2015[2] and Brauneder *et al.* 2018[6]. Updated data.

***How the data are filtered***

- Sites designated under IBA Criterion A2 (The site is known or thought to hold a significant population of at least two range-restricted species).
- All polygons, and point data with reported areas.

***How the data are processed***

- Invalid S2 geometries (see Section 1) are made valid.
- Point data buffered to reported area, `SitArea`.
- Data are unioned to remove overlap (see Section 2).

##### 8.3 IBAs under criterion A3

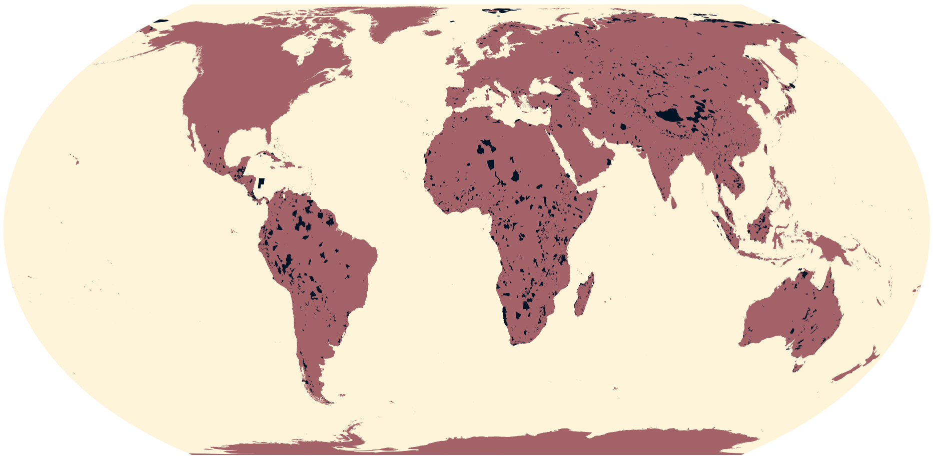

**Figure 19.** Global distribution of Important Bird and Biodiversity Areas under criterion A3.

- **Number of records**: 66 (point) and 2,984 (polygon)
- **Area (including buffered point data)**: 6,887,000 km$^2$
- **Critical Habitat type**: Potential
- **IFC PS6 Criteria triggered**: C4 (Highly Threatened or Unique Ecosystems)

***Justification and alignment with IFC PS6 criteria***
See Martin *et al.* 2015[2] and Brauneder *et al.* 2018[6]. Updated data.

***How the data are filtered***
- Sites designated under IBA Criterion A3 (The site is known or thought to hold a significant component of the group of species whose distributions are largely or wholly confined to one biome-realm).
- All polygons, and point data with reported areas.

***How the data are processed***
- Invalid S2 geometries (see Section 1) are made valid.
- Point data buffered to reported area, `SitArea`.
- Data are unioned to remove overlap (see Section 2).

#### 8.4 IBAs under criterion A4

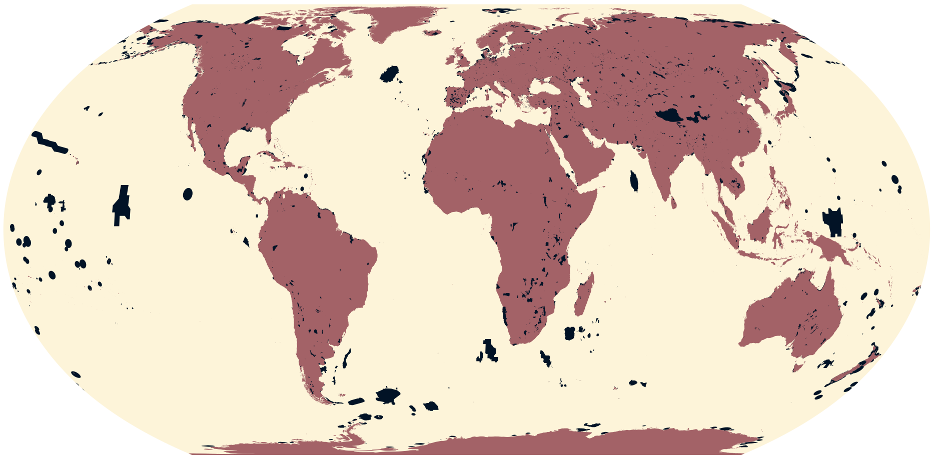

**Figure 20.** Global distribution of Important Bird and Biodiversity Areas under criterion A4.

- **Number of records**: 43 (point) and 5,615 (polygon)
- **Area (including buffered point data)**: 11,620,000 km$^2$
- **Critical Habitat type**: Likely
- **IFC PS6 Criteria triggered**: C3 (Migratory and Congregatory Species)

***Justification and alignment with IFC PS6 criteria***
See Martin *et al.* 2015[2] and Brauneder *et al.* 2018[6]. Updated data.

***How the data are filtered***
- Sites designated under IBA Criterion A4 (The site is known or thought to hold congregations of $\geq$ 1% of the global population of one or more species on a regular or predictable basis).
- All polygons, and point data with reported areas.

***How the data are processed***
- Invalid S2 geometries (see Section 1) are made valid.
- Point data buffered to reported area, `SitArea`.
- Data are unioned to remove overlap (see Section 2).

#### 8.5 IBAs under criterion B1b

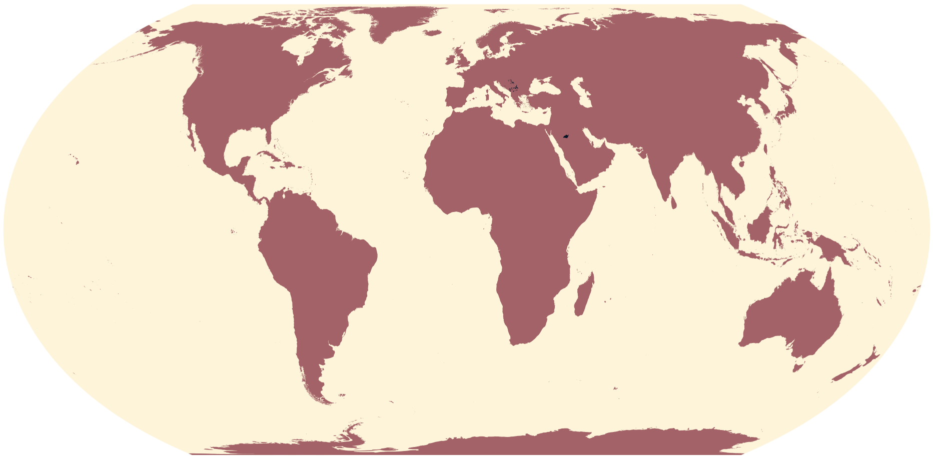

**Figure 21.** Global distribution of Important Bird and Biodiversity Areas under criterion B1b.

- **Number of records**: 1 (point) and 73 (polygon)
- **Area (including buffered point data)**: 35,390 $km^2$
- **Critical Habitat type**: Potential
- **IFC PS6 Criteria triggered**: C1 (Critically Endangered and Endangered Species)

***Justification and alignment with IFC PS6 criteria***

See Martin *et al.* 2015[2] and Brauneder *et al.* 2018[6]. Updated data.

***How the data are filtered***

- Sites designated under IBA Criterion B1b (The site is one of the 'n' most important in a country for a species with an unfavourable conservation status in the region, and for which the site-protection approach is thought to be appropriate).
- All polygons, and point data with reported areas.

***How the data are processed***

- Invalid S2 geometries (see Section 1) are made valid.
- Point data buffered to reported area, `SitArea`.
- Data are unioned to remove overlap (see Section 2).

### 9 Important Marine Mammal Sites (IMMAs)

- **Data type**: Polygon
- **Number of records**: 242
- **Original CRS**: WGS 84
- **Area - reported**: 33,330,000 $km^2$
- **Source**: IUCN-MMPATF 2023[13]

#### 9.1 IMMAs under criterion A

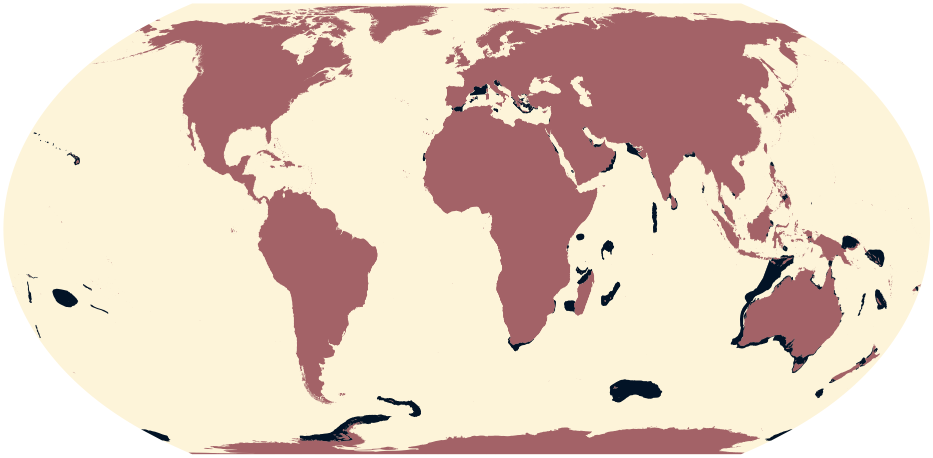

**Figure 22.** Global distribution of Important Marine Mammal Areas under criterion A.

- **Number of records**: 193
- **Area**: 8,327,000 $km^2$
- **Critical Habitat type**: Potential
- **IFC PS6 Criteria triggered**: C1 (Critically Endangered and Endangered Species)

***Justification and alignment with IFC PS6 criteria***
An Important Marine Mammal Area (IMMA) is *"a discrete portion of habitat, important for one or more marine mammal species, that has the potential to be delineated and managed for conservation"*[14]. IMMAs are identified through a science-based, expert-led three-stage process:

1. Nomination of preliminary Areas of Interest (pAoI)
2. Workshop to develop "candidate IMMAs"
3. Independent review and IMMA Status Qualification

We only consider confirmed IMMAs. Sites designated under IMMA Criterion A (Species or Population Vulnerability) align with GN72(b) and GN72(c) of GN2019[9] as they are *"Areas containing habitat important for the survival and recovery of threatened species"*, where *"threatened species"* refers to marine mammal species, subspecies or subpopulations classified as CR, EN or VU[14].

***How the data are filtered***
Sites designated under IMMA Criterion A (Species or Population Vulnerability).

***How the data are processed***
- Invalid S2 geometries (see Section 1) are made valid.
- Data are unioned to remove overlap (see Section 2).

#### 9.2 IMMAs under criterion B1

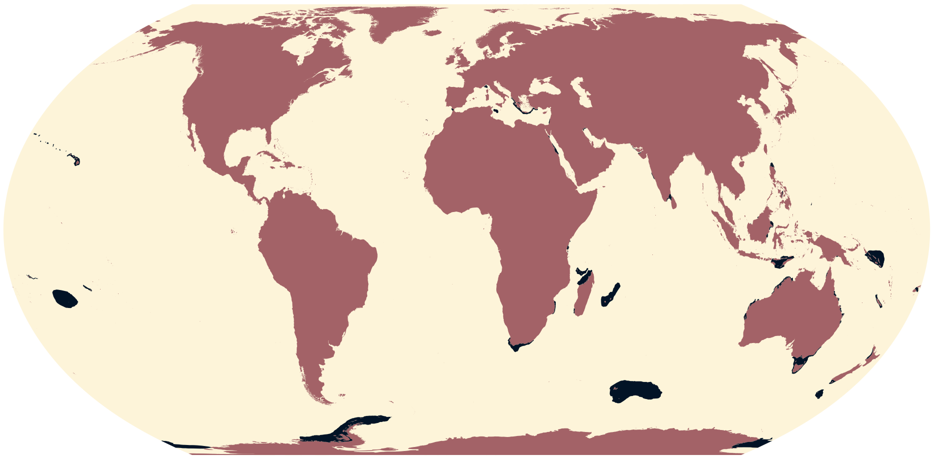

**Figure 23.** Global distribution of Important Marine Mammal Areas under criterion B1.

- **Number of records**: 106
- **Area**: 4,734,000 $km^2$
- **Critical Habitat type**: Potential
- **IFC PS6 Criteria triggered**: C2 (Endemic and Restricted-range Species)

***Justification and alignment with IFC PS6 criteria***
An Important Marine Mammal Area (IMMA) is *"a discrete portion of habitat, important for one or more marine mammal species, that has the potential to be delineated and managed for conservation"*[14]. IMMAs are identified through a science-based, expert-led three-stage process:

1. Nomination of preliminary Areas of Interest (pAoI)
2. Workshop to develop "candidate IMMAs"
3. Independent review and IMMA Status Qualification

We only consider confirmed IMMAs. Sites designated under IMMA Criterion B1 (Small and Resident Populations) align with GN75(a) of GN2019[9] as they are *"Areas supporting at least one resident population, containing an important proportion of that species or population, that are occupied consistently"*[14]. There is no definition as to what areas are considered small (cf. GN74 defining restricted-range species in marine systems as provisionally those with an extent of occupancy (EOO) of <=100,000 $km^2$), rather the size is assessed in the context of the global distribution and abundance of the species.

***How the data are filtered***
Sites designated under IMMA Criterion B1 (Small and Resident Populations).

***How the data are processed***
- Invalid S2 geometries (see Section 1) are made valid.
- Data are unioned to remove overlap (see Section 2).

#### 9.3 IMMAs under criterion B2

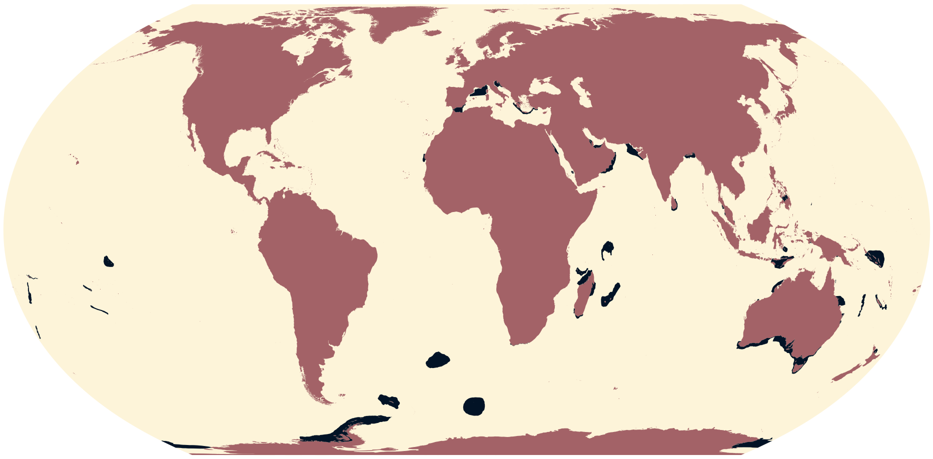

**Figure 24.** Global distribution of Important Marine Mammal Areas under criterion B2.

- **Number of records**: 106
- **Area**: 5,379,000 $km^2$
- **Critical Habitat type**: Likely
- **IFC PS6 Criteria triggered**: C3 (Migratory and Congregatory Species)

***Justification and alignment with IFC PS6 criteria***

An Important Marine Mammal Area (IMMA) is *"a discrete portion of habitat, important for one or more marine mammal species, that has the potential to be delineated and managed for conservation"*[14]. IMMAs are identified through a science-based, expert-led three-stage process:

1. Nomination of preliminary Areas of Interest (pAoI)
2. Workshop to develop "candidate IMMAs"
3. Independent review and IMMA Status Qualification

We only consider confirmed IMMAs. Sites designated under IMMA Criterion B2 (Aggregations) directly align with GN78(a) and GN78(b) of GN2019[9] as they are *"Areas with underlying qualities that support important concentrations of a species or population"*[14].

***How the data are filtered***

Sites designated under IMMA Criterion B2 (Aggregations).

***How the data are processed***

- Invalid S2 geometries (see Section 1) are made valid.
- Data are unioned to remove overlap (see Section 2).

##### 9.4 IMMAs under criteria C1, C2, and C3

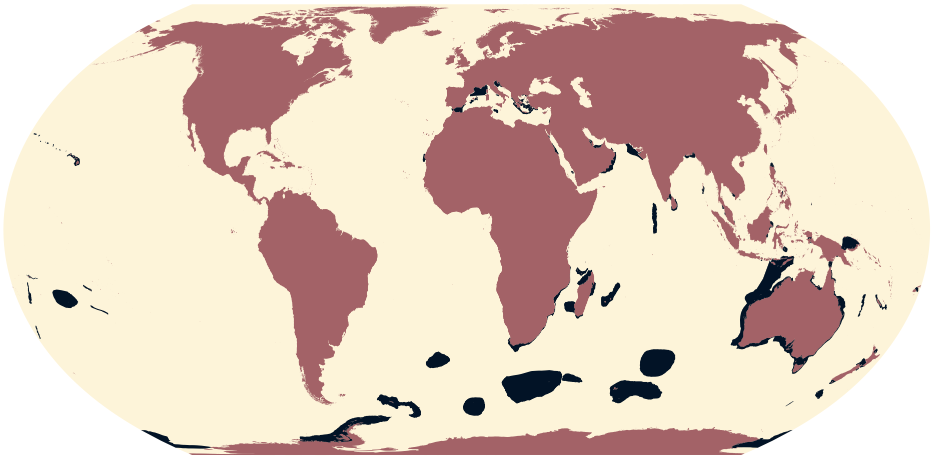

**Figure 25.** Global distribution of Important Marine Mammal Areas under criterion C.

- **Number of records**: 202
- **Area**: 12,670,000 km$^2$
- **Critical Habitat type**: Potential
- **IFC PS6 Criteria triggered**: C3 (Migratory and Congregatory Species)

***Justification and alignment with IFC PS6 criteria***

An Important Marine Mammal Area (IMMA) is *"a discrete portion of habitat, important for one or more marine mammal species, that has the potential to be delineated and managed for conservation"*[14]. IMMAs are identified through a science-based, expert-led three-stage process:

1. Nomination of preliminary Areas of Interest (pAoI)
2. Workshop to develop "candidate IMMAs"
3. Independent review and IMMA Status Qualification

We only consider confirmed IMMAs. Sites designated under IMMA Criteria C1 (Reproductive Areas), IMMA C2 (Feeding Areas), and IMMA C3 (Migration Routes) align with GN78(a) and GN78(b) of GN2019[9] as they are *"Areas that are important for a species or population to mate, give birth, and/or care for young until weaning"*[14].

***How the data are filtered***

- Sites designated under:
  - IMMA Criteria C1 (Reproductive Areas);
  - IMMA C2 (Feeding Areas); and
  - IMMA C3 (Migration Routes).

***How the data are processed***

- Invalid S2 geometries (see Section 1) are made valid.
- Data are unioned to remove overlap (see Section 2).

##### 9.5 IMMAs under criterion D1

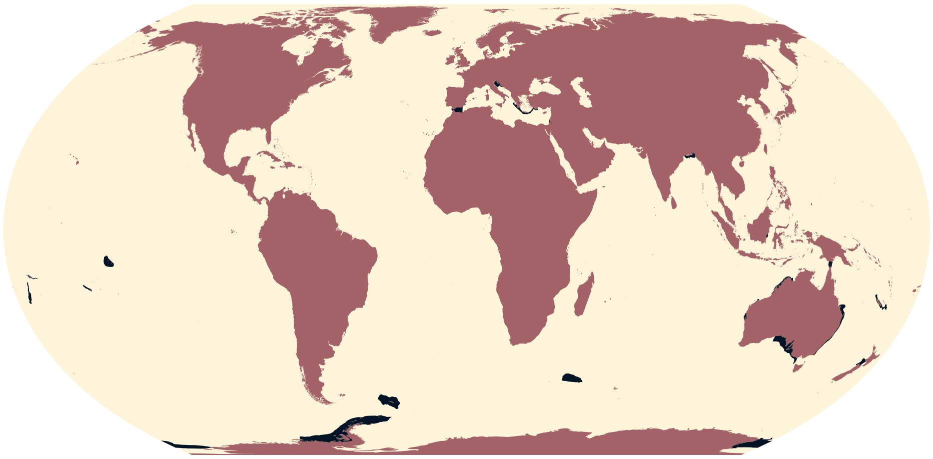

**Figure 26.** Global distribution of Important Marine Mammal Areas under criterion D1.

- **Number of records**: 56
- **Area**: 2,739,000 $km^2$
- **Critical Habitat type**: Potential
- **IFC PS6 Criteria triggered**:
  - C2 (Endemic and Restricted-range Species)
  - C4 (Highly Threatened or Unique Ecosystems)

***Justification and alignment with IFC PS6 criteria***

An Important Marine Mammal Area (IMMA) is *"a discrete portion of habitat, important for one or more marine mammal species, that has the potential to be delineated and managed for conservation"*[14]. IMMAs are identified through a science-based, expert-led three-stage process:

1. Nomination of preliminary Areas of Interest (pAoI)
2. Workshop to develop "candidate IMMAs"
3. Independent review and IMMA Status Qualification

We only consider confirmed IMMAs. Sites designated under IMMA Criterion D1 (Distinctiveness) align with GN75(a) and GN80(b) of GN2019[9] as they are *"Areas that sustain populations with important genetic, behavioural or ecologically distinctive characteristics"*[14]: GN75(a) as these sites require some form of *"geographic isolation"* and GN80(a) as example characteristics of mammal populations include *"rare or unusual behaviour or ecological linkages"*.

***How the data are filtered***

Sites designated under IMMA Criterion D1 (Distinctiveness).

***How the data are processed***

- Invalid S2 geometries (see Section 1) are made valid.
- Data are unioned to remove overlap (see Section 2).

##### 9.6 IMMAs under criterion D2

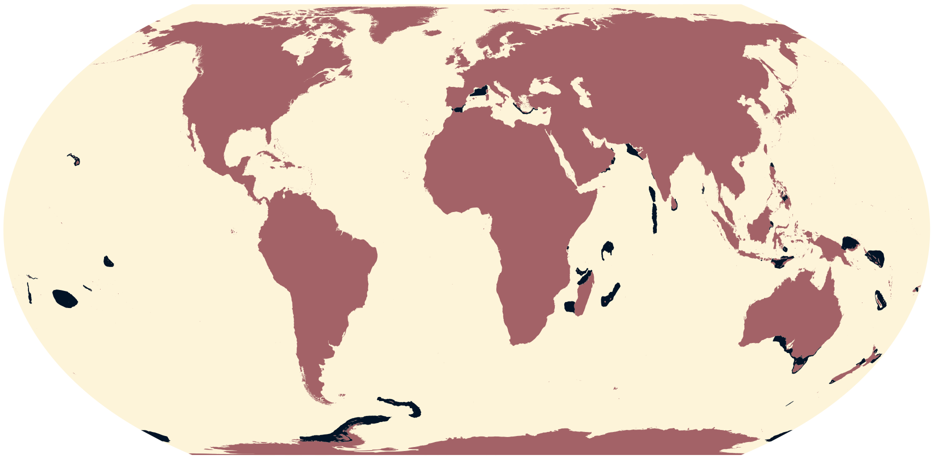

**Figure 27.** Global distribution of Important Marine Mammal Areas under criterion D2.

- **Number of records**: 77
- **Area**: 5,095,000 km$^2$
- **Critical Habitat type**: Potential
- **IFC PS6 Criteria triggered**: C4 (Highly Threatened or Unique Ecosystems)

***Justification and alignment with IFC PS6 criteria***
An Important Marine Mammal Area (IMMA) is *"a discrete portion of habitat, important for one or more marine mammal species, that has the potential to be delineated and managed for conservation"*[14]. IMMAs are identified through a science-based, expert-led three-stage process:

1. Nomination of preliminary Areas of Interest (pAoI)
2. Workshop to develop "candidate IMMAs"
3. Independent review and IMMA Status Qualification

We only consider confirmed IMMAs. Sites designated under IMMA Criterion D1 (Diversity) align with GN80(b) of GN2019[9] as they are *"Areas containing habitat that supports an important diversity of marine mammal species"*[14].

***How the data are filtered***
Sites designated under IMMA Criterion D2 (Diversity).

***How the data are processed***
- Invalid S2 geometries (see Section 1) are made valid.
- Data are unioned to remove overlap (see Section 2).

#### 10 Intact Forest Landscapes

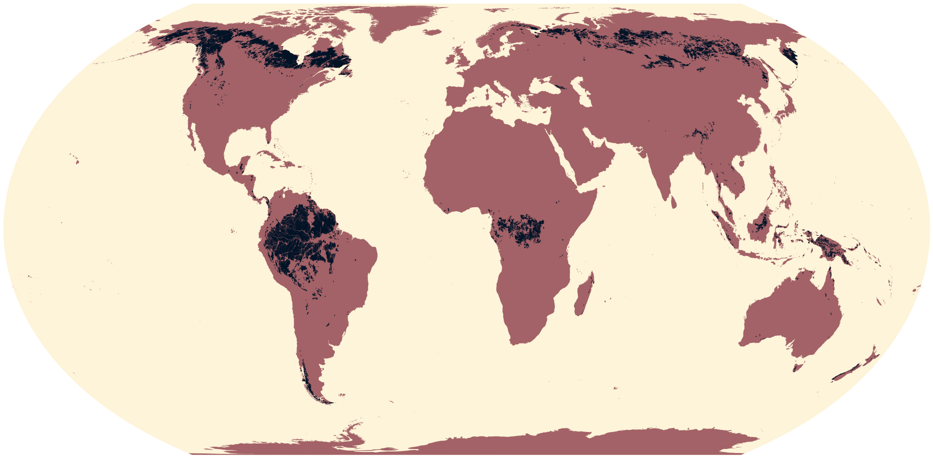

**Figure 28.** Global distribution of Intact Forest Landscapes.

- **Data type**: Polygon
- **Number of records**: 2,054
- **Original CRS**: WGS 84
- **Area**: 11,260,000 $km^2$
- **Critical habitat type**: Likely
- **IFC PS6 criteria triggered**: C4 (Highly Threatened or Unique Ecosystems)
- **Source**: Potapov *et al.* 2017[15]

***Justification and alignment with IFC PS6 criteria***

Despite being aligned with IFC PS6 criteria, Intact Forest Landscapes (IFLs) were initially excluded by Brauneder *et al.* 2018[6] on account of their spatial extent. However, on reassessment IFLs are no larger in spatial extent than other included datasets: they comprise 17.57% of Critical Habitat; IBAs under criteria A1 and A4, neither of which were previously excluded, comprise 20.76% and 17.70%. IFLs constitute 20% of all "forest zones" – areas with tree canopy cover of >20% – and are defined as *"seamless mosaics of forests and associated natural treeless ecosystems that exhibit no remotely detected signs of human activity or habitat fragmentation and are large enough to maintain all native biological diversity, including viable populations of wide-ranging species"*[15]. Intactness correlates well with the full conservation value of forest landscapes[16,17,18]. For this reason, they are likely to be of high priority for conservation and therefore align directly with GN80(b).

***How the data are filtered***

No filtering of data from source.

***How the data are processed***

- Invalid S2 geometries (see Section 1) are made valid.
- Data are unioned to remove overlap (see Section 2).

#### 11 Irreplaceable protected areas

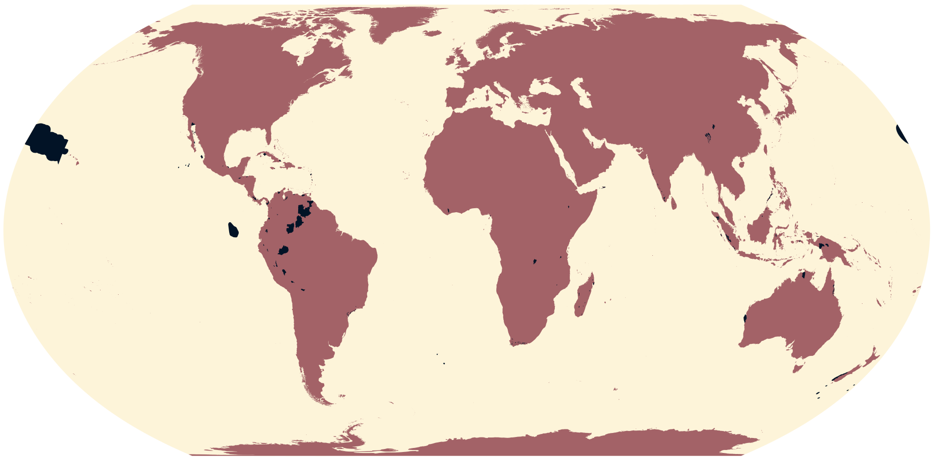

**Figure 29.** Global distribution of irreplaceable protected areas.

- **Data type**: Polygon
- **Number of records**: 115
- **Original CRS**: WGS 84
- **Area**: 2,484,000 km$^2$
- **Critical Habitat type**: Likely
- **IFC PS6 Criteria triggered**: C4 (Highly Threatened or Unique Ecosystems)
- **Source**: Le Saout *et al.* 2013[19]

***Justification and alignment with IFC PS6 criteria***
See Brauneder *et al.* 2018[6]. No change to data.

***How the data are filtered***
No filtering of data from source.

***How the data are processed***
- Invalid S2 geometries (see Section 1) are made valid.
- Data are unioned to remove overlap (see Section 2).

#### 12 IUCN Red List of Threatened Species

- **Data type**: Polygon (we exclude point data)
- **Number of records**: 137,500
- **Original CRS**: WGS 84
- **Source**: IUCN 2024[20]
- **Version**: 2024_1

***How the data are filtered***

- Critically Endangered (CR), Endangered (EN) or Vulnerable (VU) species classified under Criterion D for CR or EN (Population size estimated to number fewer than 50(CR)/250(EN) mature individuals) and D2 for VU (Population with a very restricted area of occupancy (typically less than 20 $km^2$) or number of locations (typically five or fewer) such that it is prone to the effects of human activities or stochastic events within a very short time period in an uncertain future, and is thus capable of becoming CR or even Extinct in a very short time period).
- Exclude marine ranges (`biome_marine==false`).
- Species in Family *Hominidae*.
- Exclude *Extinct (post 1500)* or *Probably Extant* (deprecated category) ranges.
- Ranges with areas ≥ three times the standard deviation of the data (see Technical Validation in main text). This exercise removes areas disproportionately large. Furthermore, some of the records removed include the northern white rhino, of which there is only one individual left, and the ivory-billed woodpecker, which is argued to be extinct.

***How the data are processed***

Invalid S2 geometries (see Section 1) are made valid.

##### 12.1 Critically Endangered species under criterion D

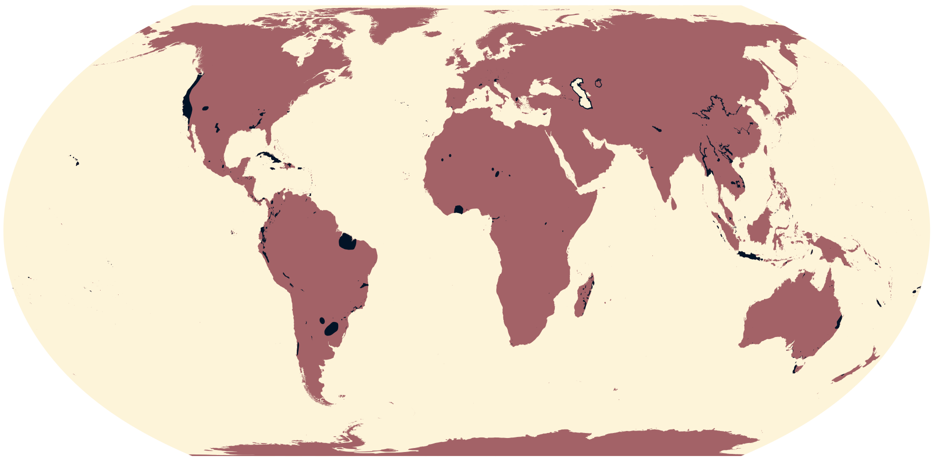

**Figure 30.** Global distribution of Critically Endangered species under criterion D.

- **Number of records**: 973
- **Area**: 2,759,000 $km^2$
- **Critical Habitat type**: Likely
- **IFC PS6 Criteria triggered**: C1 (Critically Endangered or Endangered Species)

***Justification and alignment with IFC PS6 criteria***

See Brauneder *et al.* 2018[6]. Updated data.

***How the data are filtered***

No further filtering of data.

***How the data are processed***

- Invalid S2 geometries (see Section 1) are made valid.
- Data are unioned to remove overlap (see Section 2).

#### 12.2 Endangered species under criterion D

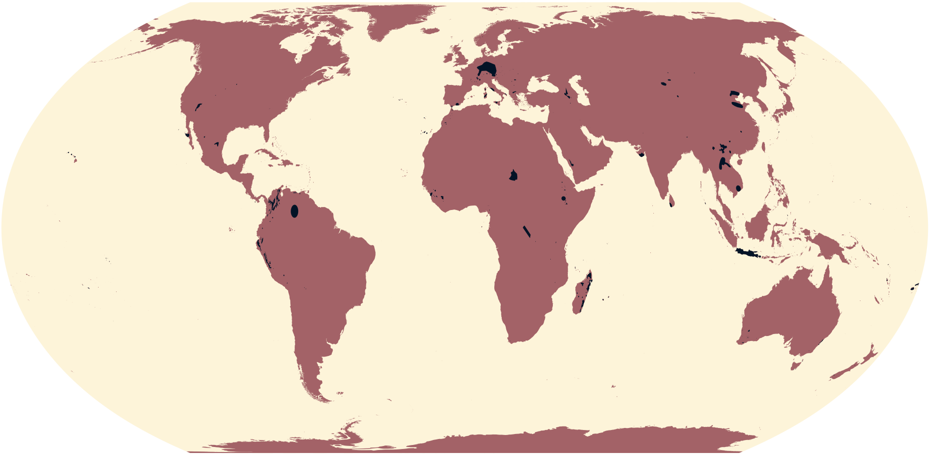

**Figure 31.** Global distribution of Endangered species under criterion D.

- **Number of records**: 346
- **Area**: 1,417,000 km$^2$
- **Critical Habitat type**: Likely
- **IFC PS6 Criteria triggered**: C1 (Critically Endangered or Endangered Species)

***Justification and alignment with IFC PS6 criteria***
See Brauneder *et al.* 2018[6]. Updated data.

***How the data are filtered***
No further filtering of data from source.

***How the data are processed***
- Invalid S2 geometries (see Section 1) are made valid.
- Data are unioned to remove overlap (see Section 2).

##### 12.3 Vulnerable species under criterion D2

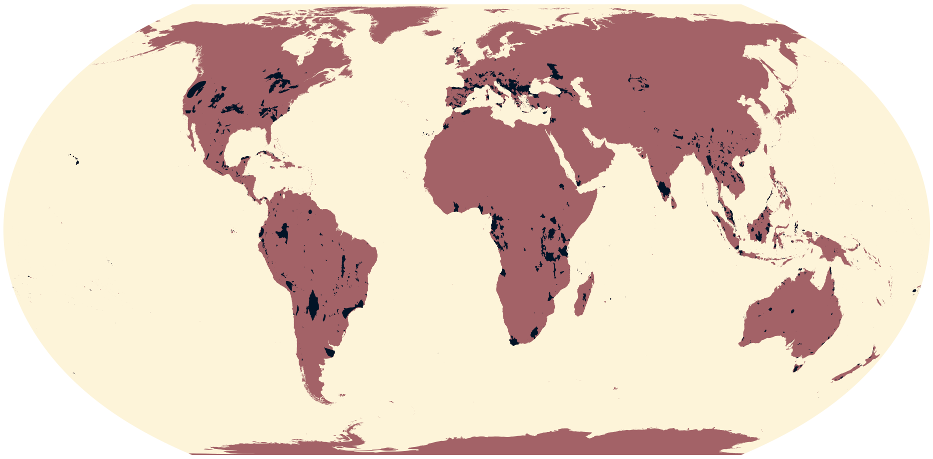

**Figure 32.** Global distribution of Vulnerable species under criterion D2.

- **Number of records**: 2,217
- **Area**: 6,097,000 $km^2$
- **Critical Habitat type**: Potential
- **IFC PS6 Criteria triggered**: C1 (Critically Endangered or Endangered Species)

***Justification and alignment with IFC PS6 criteria***
Species classified as VU under Criterion D2 represent a *"Population with a very restricted area of occupancy (typically less than 20 $km^2$) or number of locations (typically five or fewer) such that it is prone to the effects of human activities or stochastic events within a very short time period in an uncertain future, and is thus capable of becoming CR or even Extinct in a very short time period"*. This aligns with GN72(b), although whether species classified in this manner also meet the thresholds of GN72(a), as required, is uncertain.

***How the data are filtered***
No further filtering of data.

***How the data are processed***
- Invalid S2 geometries (see Section 1) are made valid.
- Data are unioned to remove overlap (see Section 2).

#### 13 Great apes habitat

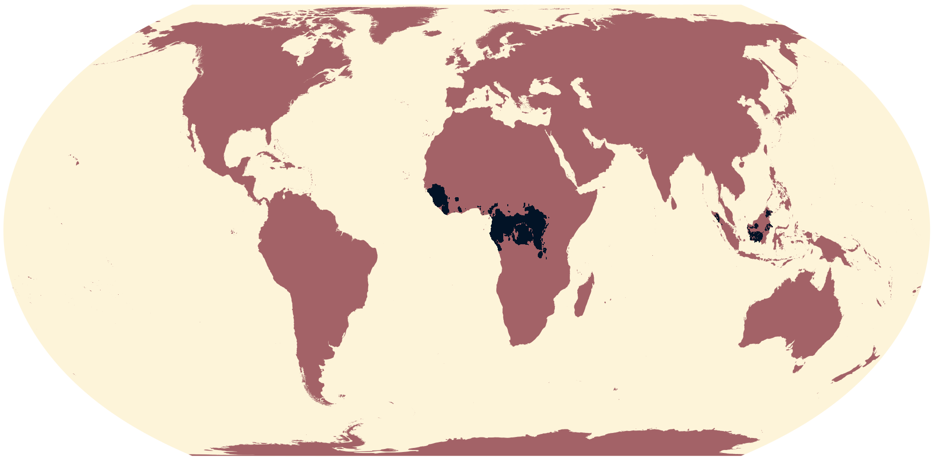

**Figure 33.** Global distribution of great apes.

- **Data type**: Raster
- **Original CRS**: WGS 84
- **Original resolution**: 100m
- **Extent**: Global
- **Gridded area of feature at 1-km resolution**: 3,016,000
- **Critical Habitat type**: Likely
- **IFC PS6 Criteria triggered**: C1 (Critically Endangered or Endangered Species)
- **Source**: Lumbierres *et al.* 2022[21]

***Justification and alignment with IFC PS6 criteria***

GN73 of GN2019 requests *"special consideration"* for non-human great ape species. Any area with great apes present is likely to be treated as Critical Habitat. Lumbierres *et al.* 2022[21] produced Area of Habitat (AoH) – *"the habitat available to a species, that is, habitat within its range"* – maps for 5,481 terrestrial mammal species, including great apes, at ~100m resolution. AoH maps are produced by subtracting unsuitable habitat from a species' geographic range and therefore indicate a high probability of presence on the ground. As such, great apes habitat is classified as Likely Critical Habitat.

***How the data are processed***

- Area of habitat data for each of the non-human great apes species at ~100m resolution are aggregated to 30 arcseconds resolution using a modal function.
- Data are then merged together to form a data layer at global extent for all great ape species.

#### 14 Mangroves

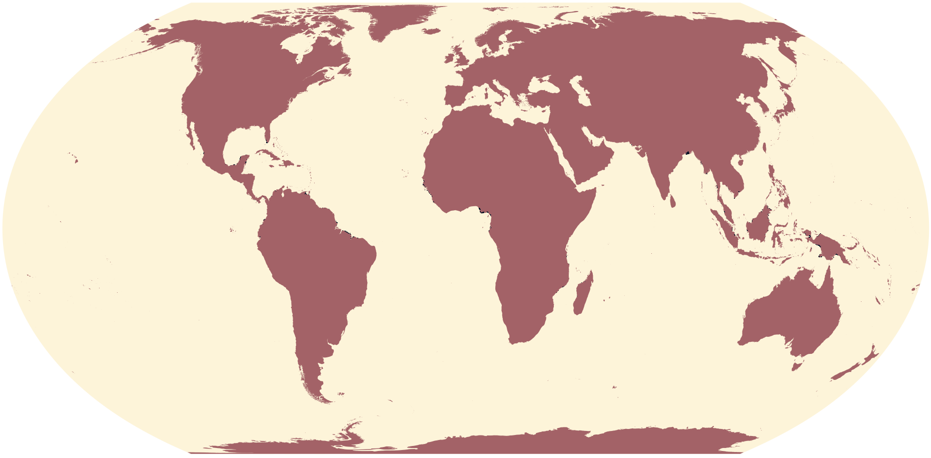

**Figure 34.** Global distribution of mangroves.

- **Data type**: Raster
- **Original CRS**: WGS 84
- **Original resolution**: 0.0002222
- **Extent**: -180-39.0118033
- **Gridded area of feature at 1-km resolution**: 115,000
- **Critical Habitat type**: Likely
- **IFC PS6 Criteria triggered**: C4 (Highly Threatened or Unique Ecosystems)
- **Source**: Bunting *et al.* 2022[22]

***Justification and alignment with IFC PS6 criteria***
See Martin *et al.* 2015[2] and Brauneder *et al.* 2018[6]. Updated data.

***How the data are processed***
- Raster tiles are mosaicked to a global extent.
- Raster mosaic is then aggregated to a 30 arcseconds resolution using the following function: `function(x,...){sum(x,na.rm` that returns the percentage of cells identified as mangrove.
- Resulting raster is then resampled to a template global raster.
- Cells are then classified as a presence when the percentage is above a certain threshold ($\geq 50\%$ in this case - see Figure 35 for a sensitivity analysis of this).

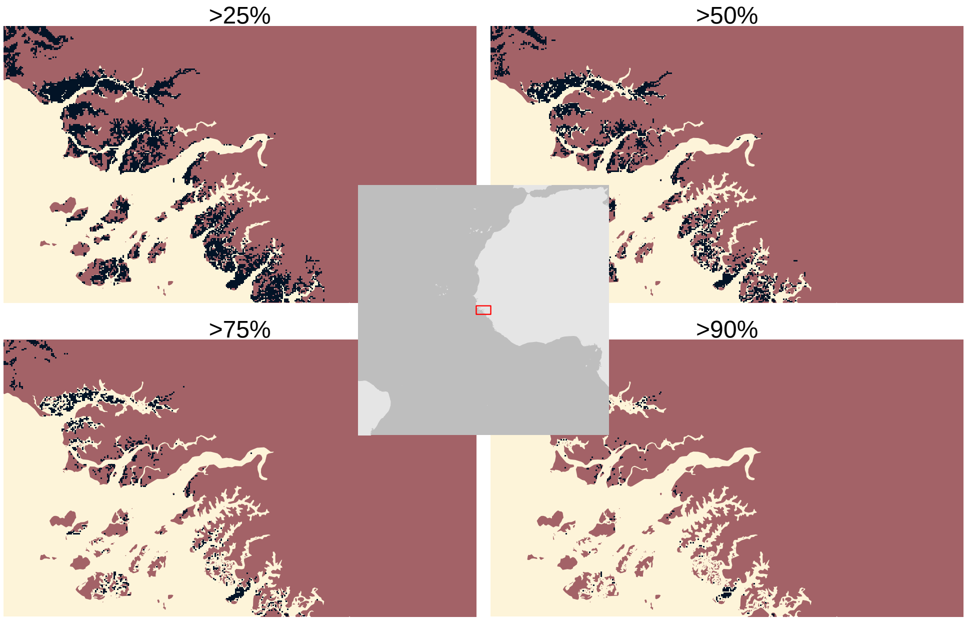


**Figure 35.** Sensitivity analysis of % presence cutoff values. Plots show effect of setting the % presence to >25, >50, >75, >90% on the output binary distribution. We select 50% for use in this analysis.

#### 15 Saltmarsh

- **Data type**: Point and polygon
- **Number of records**: 2,946
- **Original CRS**: WGS 84
- **Area (polygons)**: 54,540 km$^2$
- **Critical Habitat type**: Likely
- **IFC PS6 Criteria triggered**: C4 (Highly Threatened or Unique Ecosystems)
- **Source**: Mcowen *et al.* 2017[23]

***Justification and alignment with IFC PS6 criteria***

See Martin *et al.* 2015[2] and Brauneder *et al.* 2018[6]. No change to data.

***How the data are filtered***

No filtering of data from source.

***How the data are processed***

- Invalid S2 geometries (see Section 1) are made valid.
- Data are unioned to remove overlap (polygons) and duplicates (points) (see Section 2).

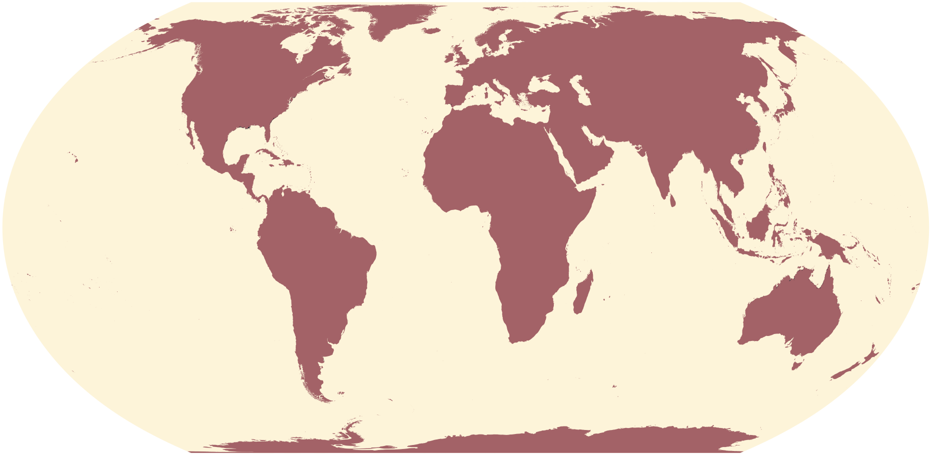

**Figure 36.** Global distribution of saltmarshes.

#### 16 Sea turtle nesting sites

- **Number of records**: 5,581
- **Original CRS**: WGS 84
- **Data type**: Point
- **Source**: Halpin *et al.* 2009; Kot *et al.* 2020[24,25]

##### 16.1 All sea turtle species

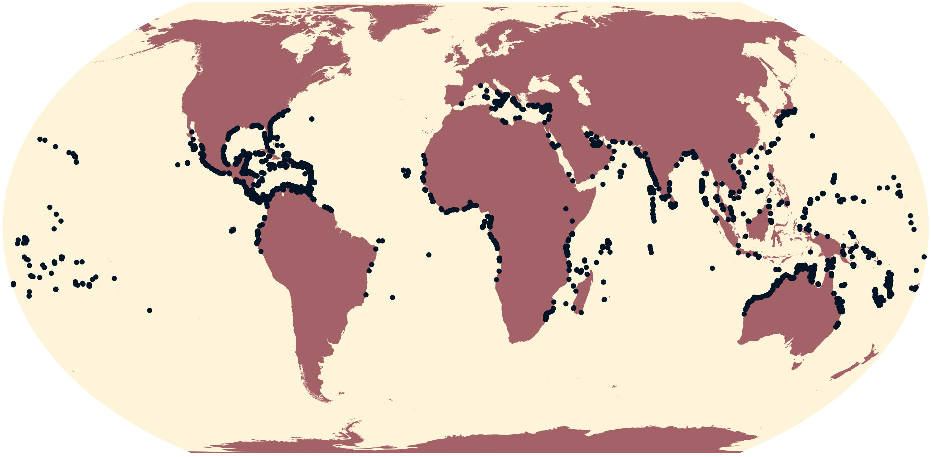

**Figure 37.** Global distribution of all sea turtle species nesting sites.

- **Number of records**: 5,581
- **Data type**: Point
- **Critical habitat type**: Potential
- **IFC PS6 criteria triggered**:
  - C3 (Migratory and Congregatory Species)
  - C4 (Highly Threatened or Unique Ecosystems)
- **Source**:

***Justification and alignment with IFC PS6 criteria***
See Martin *et al.* 2015[2] and Brauneder *et al.* 2018[6]. Updated data.

***How the data are filtered***
No filtering of data from source.

***How the data are processed***
Data are unioned to remove duplicates (see Section 2).

##### 16.2 Critically Endangered and Endangered sea turtle species

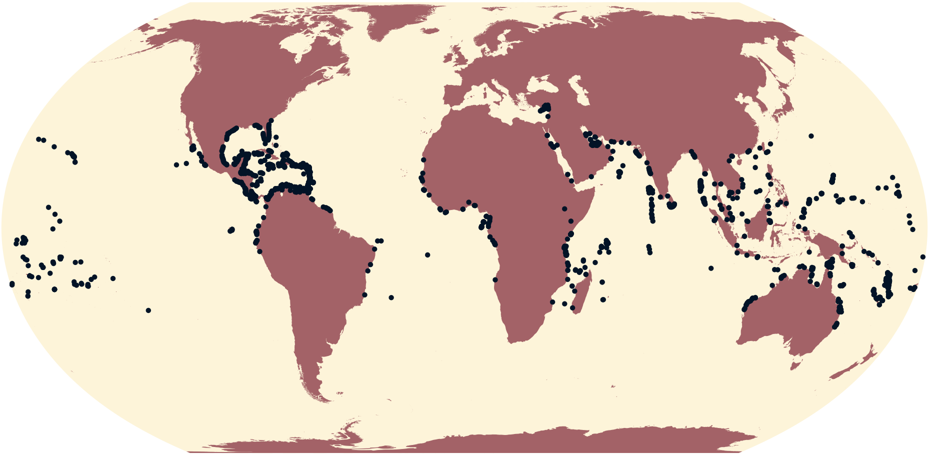

**Figure 38.** Global distribution of Critically Endangered or Endangered sea turtle species nesting sites.

- **Number of records**: 3,250
- **Data type**: Point
- **Critical habitat type**: Likely
- **IFC PS6 criteria triggered**: C1 (Critically Endangered and Endangered Species)

***Justification and alignment with IFC PS6 criteria***
See Martin *et al.* 2015[2] and Brauneder *et al.* 2018[6]. Updated data.

***How the data are filtered***
Records identified as belonging to Critically Endangered or Endangered sea turtles: Green Sea Turtle, Hawksbill Sea Turtle, and Kemp's Ridley.

***How the data are processed***
Data are unioned to remove duplicates (see Section 2).

#### 17 Seagrass beds

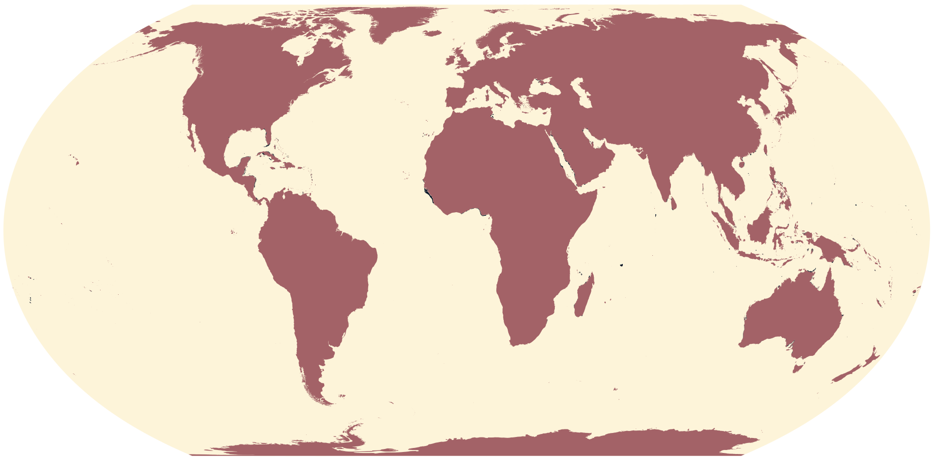

**Figure 39.** Global distribution of seagrasses.

- **Data type**: Point and polygon
- **Number of records**: 310,800
- **Original CRS**: WGS84
- **Area (polygons)**: 314,900 $km^2$
- **Critical habitat type**: Likely
- **IFC PS6 criteria triggered**: C4 (Highly Threatened or Unique Ecosystems)
- **Source**: UNEP-WCMC 2005[26]

***Justification and alignment with IFC PS6 criteria***
See Martin *et al.* 2015[2]. No change to data.

***How the data are filtered***
No filtering of data from source.

***How the data are processed***
- Invalid S2 geometries (see Section 1) are made valid.
- Data are unioned to remove overlap (polygons) and duplicates (points) (see Section 2).

#### 18 Seamounts

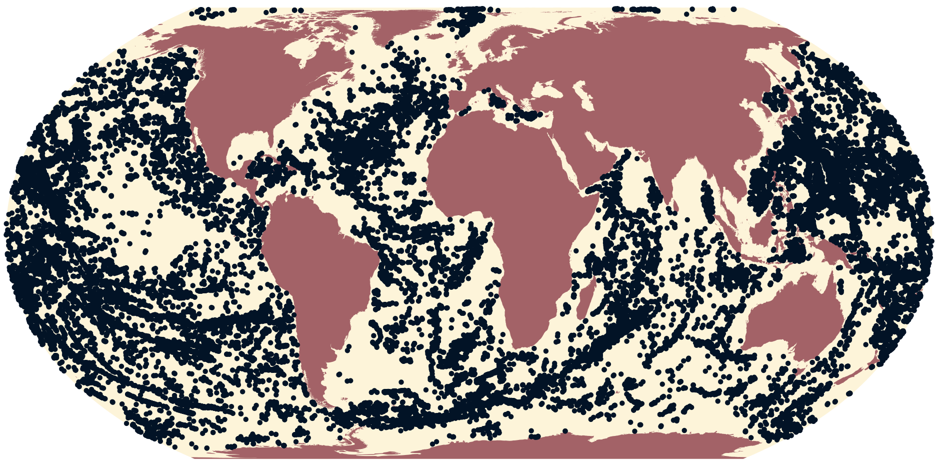

**Figure 40.** Global distribution of seamounts.

- **Data type**: Point
- **Number of records**: 33,450
- **Original CRS**: WGS 84
- **Critical habitat type**: Potential
- **IFC PS6 criteria triggered**: C4 (Highly Threatened or Unique Ecosystems)
- **Source**: Yesson *et al.* 2011[27]

***Justification and alignment with IFC PS6 criteria***
See Martin *et al.* 2015[2]. No change to data.

***How the data are filtered***
No filtering of data from source.

***How the data are processed***
- Invalid S2 geometries (see Section 1) are made valid.
- Data are unioned to remove duplicates (see Section 2).

#### 19 Tiger Conservation Landscapes

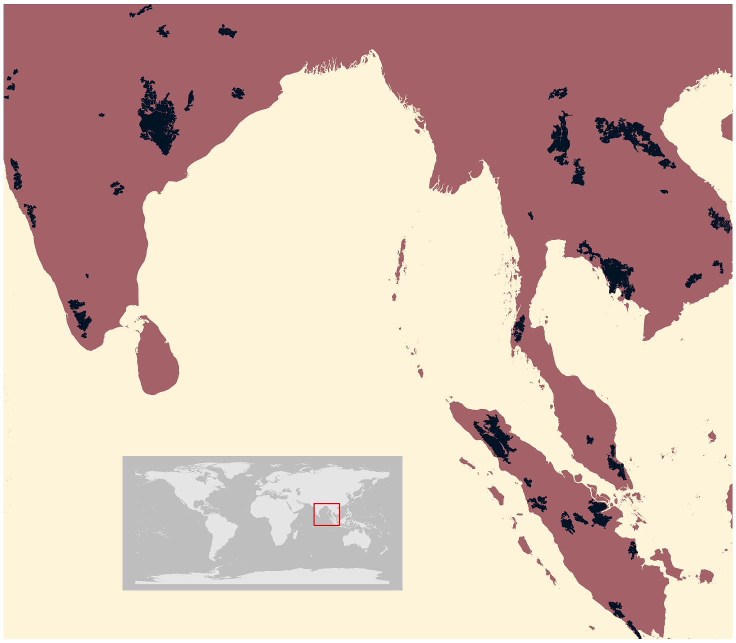

**Figure 41.** Global distribution of Tiger Conservation Landscapes (inset: South Asia).

- **Data type**: Polygon
- **Number of records**: 47
- **Original CRS**: WGS 84
- **Critical habitat type**: Likely
- **IFC PS6 criteria triggered**: C1 (Critically Endangered and Endangered Species)
- **Source**: Sanderson *et al.* 2023[28]

***Justification and alignment with IFC PS6 criteria***

See Brauneder *et al.* 2018[6]. Updated data.

***How the data are filtered***

Current tiger habitat only: `tx2_tcl==0`.

***How the data are processed***

- Invalid S2 geometries (see Section 1) are made valid.
- Data are unioned to remove overlap (see Section 2).

#### 20 Tropical dry forest

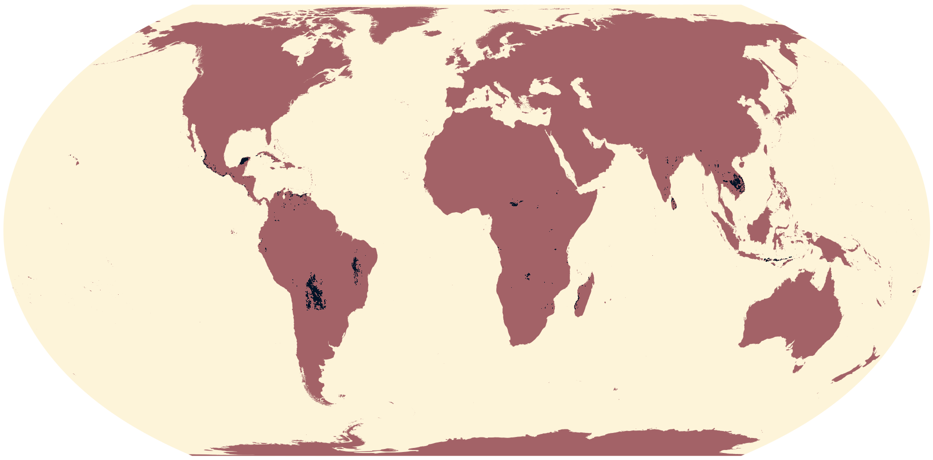

**Figure 42.** Global distribution of tropical dry forests.

- **Data type**: Raster
- **Original CRS**: WGS 84
- **Original resolution**: 0.004497
- **Extent**: -180-56.4518079.98
- **Gridded area of feature at 1-km resolution**: 840,800
- **Critical habitat type**: Potential
- **IFC PS6 criteria triggered**: C4 (Highly Threatened or Unique Ecosystems)
- **Source**: Miles *et al.* 2006[29]

***Justification and alignment with IFC PS6 criteria***
See Brauneder *et al.* 2018[6]. No change to data.

***How the data are processed***
- Missing data are reclassified to 0.
- Raster aggregated from native resolution to 30 arcseconds using the mode of neighbouring cells before resampling to template raster using nearest neighbour.

#### 21 Tropical moist forest

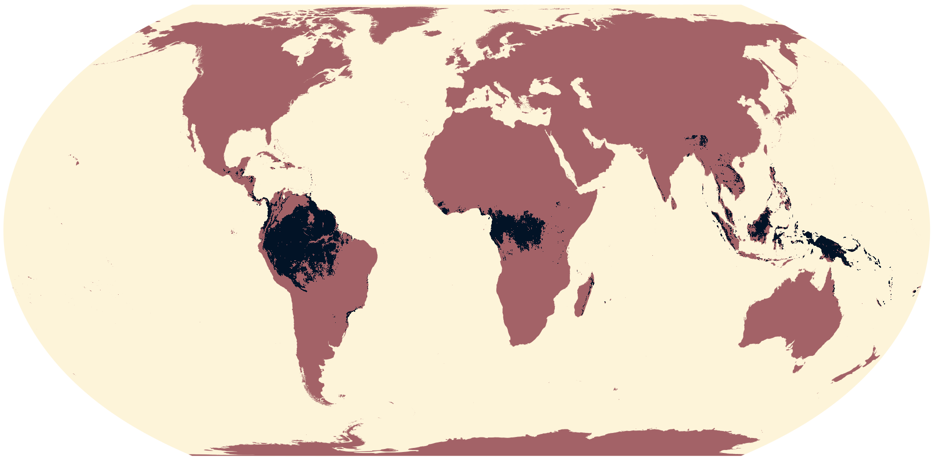

**Figure 43.** Global distribution of moist tropical forests.

- **Data type**: Raster
- **Original CRS**: WGS 84
- **Original resolution**: 0.0002695
- **Extent**: -110-40.1218030.09
- **Gridded area of feature at 1-km resolution**: 9,037,000
- **Critical habitat type**: Likely
- **IFC PS6 criteria triggered**: C4 (Highly Threatened or Unique Ecosystems)
- **Source**: Vancutsem *et al.* 2021[30]

***How the data are processed***

- Raster tiles are mosaicked to a global extent.
- Raster mosaic is then aggregated to a 30 arcseconds resolution using the following function: `function(x,...){sum((x == 10),na.rm = T)/length(x)}` that returns the percentage of cells equaling 10, "Undisturbed tropical moist forest".
- Resulting raster is then resampled to a template global raster.
- Cells are then classified as a presence when the percentage is above a certain threshold ($\geq$ 50% in this case - see Figure 44 for a sensitivity analysis of this).

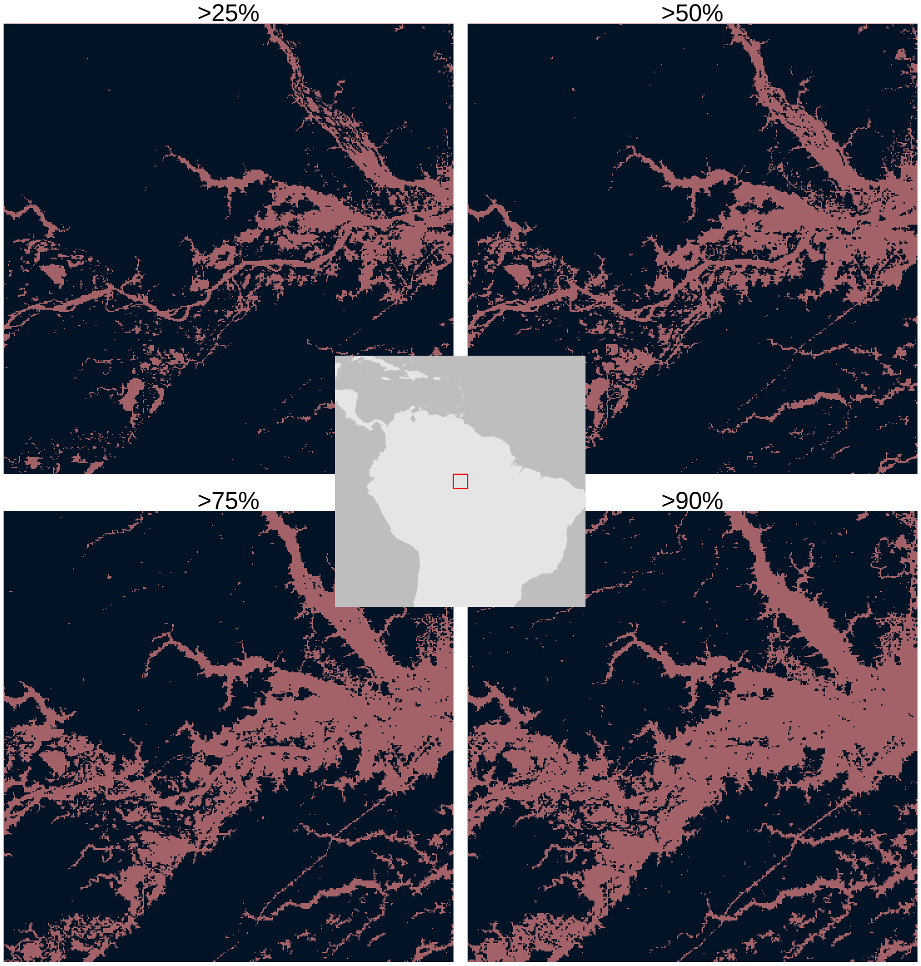


**Figure 44.** Sensitivity analysis of % presence cutoff values.

#### 22 Tropical montane cloud forests

- **Data type**: Raster
- **Original CRS**: Equirectangular
- **Original resolution**: 1,000
- **Extent**: -20,040,000-10,020,00020,040,00010,020,000
- **Gridded area of feature at 1-km resolution**: 1,567,000
- **Critical habitat type**: Likely
- **IFC PS6 criteria triggered**: C4 (Highly Threatened or Unique Ecosystems)
- **Source**: Karger *et al.* 2021[31]

***Justification and alignment with IFC PS6 criteria***

See Brauneder *et al.* 2018[6]. New data source.

***How the data are filtered***

No filtering of data from source.

***How the data are processed***

Data are unioned to remove duplicates (see Section 2).

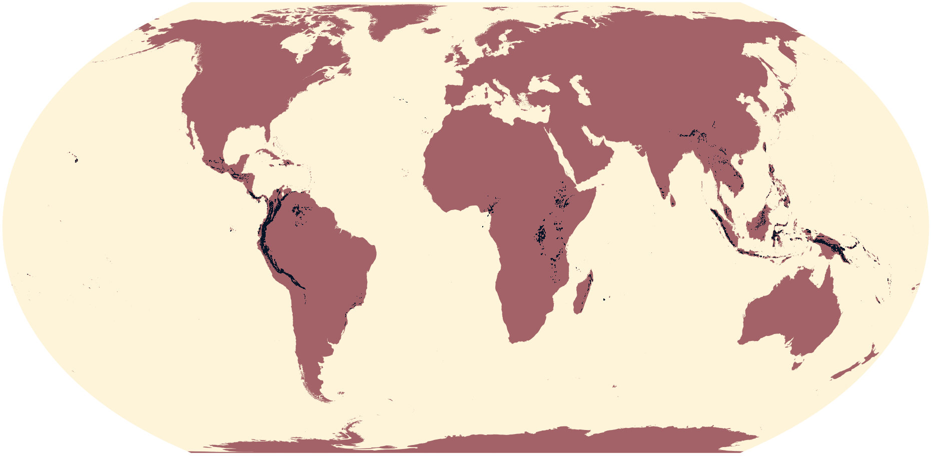

**Figure 45.** Global distribution of tropical montane cloud forests.

#### 23 Warm water coral reefs

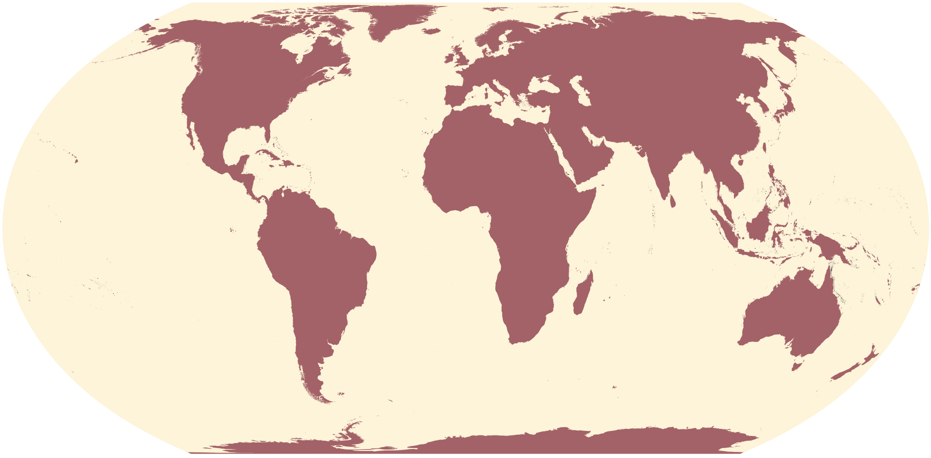

**Figure 46.** Global distribution of warm water coral reefs.

- **Data type**: Point and polygon
- **Number of records**: 18,430
- **Original CRS**: WGS 84
- **Area (polygons)**: 149,600
- **Critical habitat type**: Likely

- **IFC PS6 criteria triggered**:
  - C4 (Highly Threatened or Unique Ecosystems)
  - C5 (Key Evolutionary Processes)
- **Source**: UNEP-WCMC, WorldFish, World Resources Institute, and The Nature Conservancy 2010[32]

***Justification and alignment with IFC PS6 criteria***
See Martin *et al.* 2015[2]. No change to data.

***How the data are filtered***
No filtering of data from source.

***How the data are processed***
- Invalid S2 geometries (see Section 1) are made valid.
- Data are unioned to remove overlap (polygons) and duplicates (point) (see Section 2).

#### 24 World Database on Protected Areas (WDPA)

- **Data type**: Point and polygon
- **Number of records**: 293,300 (polygons); 11,940 (points)
- **Original CRS**: WGS 84
- **Area (polygons - reported)**: 88,290,000 km$^2$
- **Area (point - reported)**: 3,684,000 km$^2$
- **Source**: UNEP-WCMC and IUCN 2024[33]
- **Version**: October 2024

For all data sourced from the WDPA, point data were buffered to reported area, as recommended by the WDPA Manual v1.6.

##### 24.1 All Ramsar sites

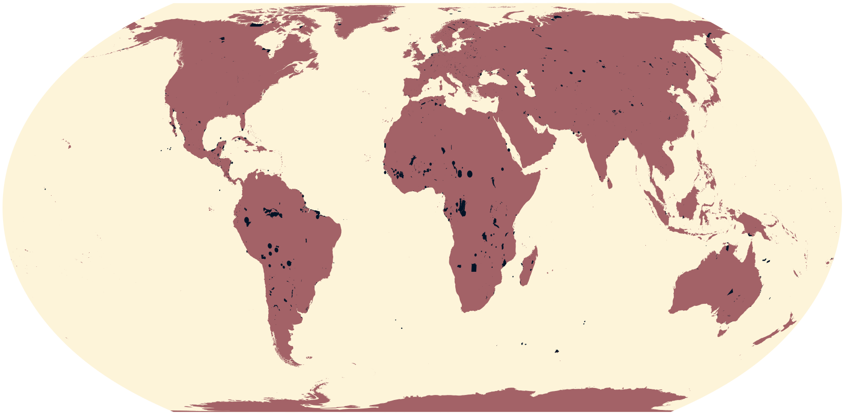

**Figure 47.** Global distribution of all Ramsar sites.

- **Number of records**: 1,919 (polygons); 430 (points)
- **Area**: 2,072,000 km$^2$
- **Critical habitat type**: Likely
- **IFC PS6 criteria triggered**: C4 (Highly Threatened or Unique Ecosystems)

***Justification and alignment with IFC PS6 criteria***
See Martin *et al.* 2015[2] and Brauneder *et al.* 2018[6]. Updated data.

***How the data are filtered***

- Sites where Status is not "Proposed" or "Not Reported".
- Sites labelled as "Ramsar Site, Wetland of International Importance", with Designation Type "International".
- Point data with present, non-zero reported area (`REP_AREA > 0`).

***How the data are processed***

- Invalid S2 geometries (see Section 1) are made valid.
- Point data buffered to reported area, `REP_AREA`.
- Data are unioned to remove overlap (see Section 2).

##### 24.2 Ramsar sites under criterion 2

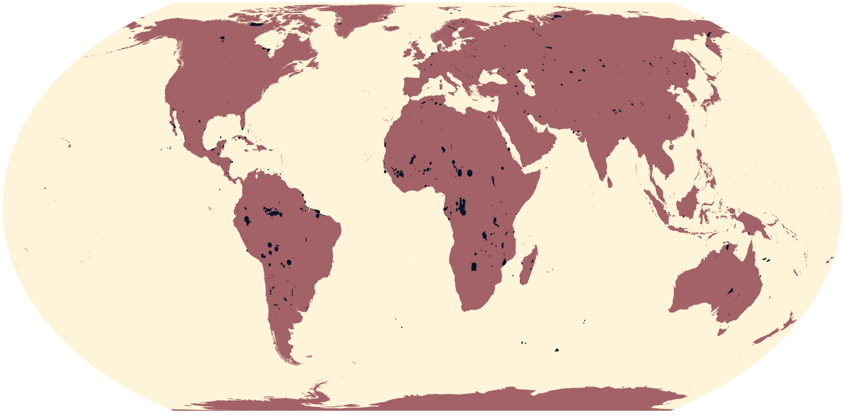

**Figure 48.** Global distribution of all Ramsar sites under criterion 2.

- **Number of records**: 1,373 (polygons); 357 (points)
- **Area**: 2,072,000 $km^2$
- **Critical habitat type**: Likely
- **IFC PS6 criteria triggered**: C1 (Critically Endangered and Endangered Species)

***Justification and alignment with IFC PS6 criteria***
See Martin *et al.* 2015[2] and Brauneder *et al.* 2018[6]. Updated data.

***How the data are filtered***
- Sites where status is not "Proposed" or "Not Reported".
- Sites labelled as "Ramsar Site, Wetland of International Importance", with Designation Type "International"
- Sites designated under Ramsar Sites Criterion 2 (Sites supporting vulnerable, endangered, or critically endangered species or threatened ecological communities).
- All polygon data and point data with present, non-zero reported area (`REP_AREA > 0`).

***How the data are processed***
- Invalid S2 geometries (see Section 1) are made valid.
- Point data buffered to reported area, `REP_AREA`.
- Data are unioned to remove overlap (see Section 2).

##### 24.3 Ramsar sites under criteria 5 and 6

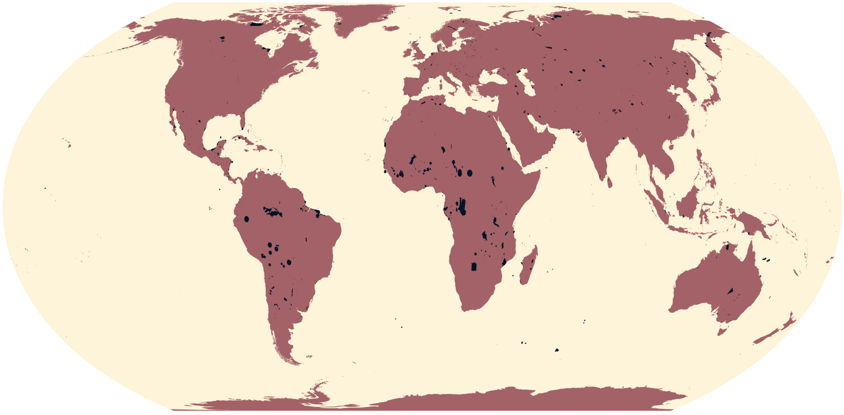

**Figure 49.** Global distribution of all Ramsar sites under criteria 5 and 6.

- **Number of records**: 838 (polygons); 191 (points)
- **Area**: 1,979,000 km$^2$
- **Critical habitat type**: Likely
- **IFC PS6 criteria triggered**: C3 (Migratory and Congregatory Species)

***Justification and alignment with IFC PS6 criteria***

See Martin *et al.* 2015[2] and Brauneder *et al.* 2018[6]. Updated data.

***How the data are filtered***

- Sites where status is not "Proposed" or "Not Reported".
- Sites labelled as "Ramsar Site, Wetland of International Importance", with Designation Type "International"
- Sites designated under:
  - Ramsar Sites Criteria 5 (Sites that regularly support 20,000 or more waterbirds); and
  - Ramsar Sites Criteria 6 (Sites that regularly support 1% of the individuals in a population of one species or subspecies of waterbird).
- All polygon data and point data with present, non-zero reported area (`REP_AREA > 0`).

***How the data are processed***

- Invalid S2 geometries (see Section 1) are made valid.
- Point data buffered to reported area, `REP_AREA`.
- Data are unioned to remove overlap (see Section 2).

##### 24.4 Ramsar sites under criteria 1 and 3

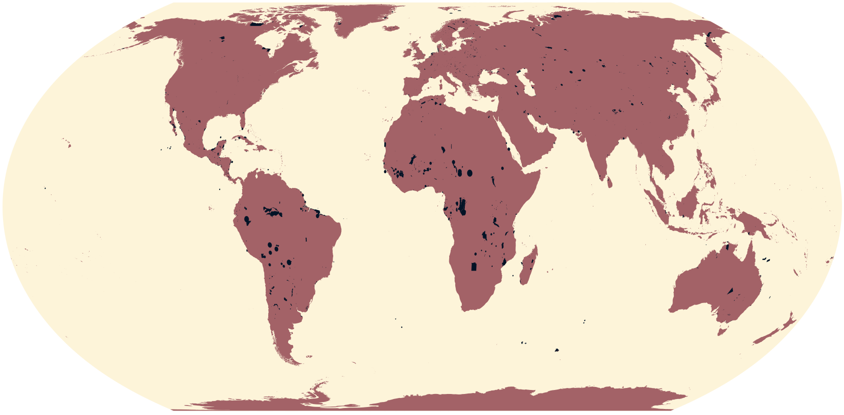

**Figure 50.** Global distribution of all Ramsar sites under criteria 1 and 3.

- **Number of records**: 1,484 (polygons); 364 (points)
- **Area**: 2,072,000 km$^2$
- **Critical habitat type**: Likely
- **IFC PS6 criteria triggered**: C4 (Highly Threatened or Unique Ecosystems)

***Justification and alignment with IFC PS6 criteria***
See Martin *et al.* 2015[2] and Brauneder *et al.* 2018[6]. Updated data.

***How the data are filtered***
- Sites where status is not "Proposed" or "Not Reported".
- Sites labelled as "Ramsar Site, Wetland of International Importance", with Designation Type "International"
- Sites designated under:
  - Ramsar Sites Criteria 1 (Sites that contain a representative, rare, or unique example of a natural or near-natural wetland type found within the appropriate biogeographic region); and
  - Ramsar Sites Criteria 3 (Sites that support populations of plant and/or animal species important for maintaining the biological diversity of a particular biogeographic region).
- All polygon data and point data with present, non-zero reported area (`REP_AREA > 0`).

***How the data are processed***
- Invalid S2 geometries (see Section 1) are made valid.
- Point data buffered to reported area, `REP_AREA`.
- Data are unioned to remove overlap (see Section 2).

##### 24.5 Ramsar sites under criteria 4, 7, 8 and 9

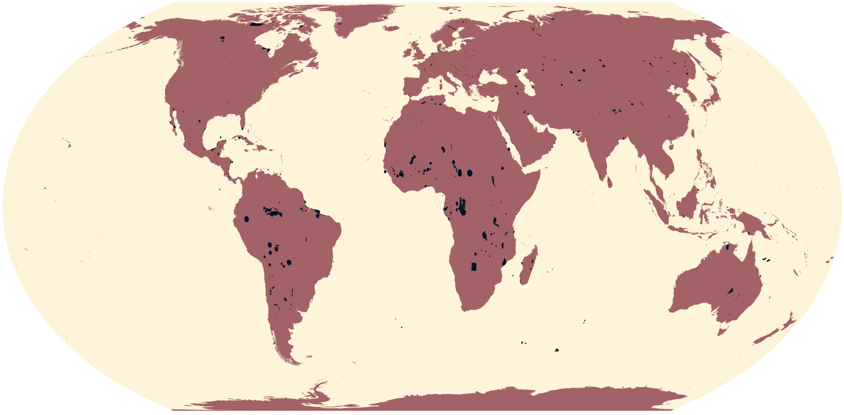

**Figure 51.** Global distribution of all Ramsar sites under criteria 4, 7, 8 and 9.

- **Number of records**: 1,174 (polygons); 322 (points)
- **Area**: 1,894,000 km$^2$
- **Critical habitat type**: Potential
- **IFC PS6 criteria triggered**: C3 (Migratory and Congregatory Species)

***Justification and alignment with IFC PS6 criteria***
See Martin *et al.* 2015[2] and Brauneder *et al.* 2018[6]. Updated data.

***How the data are filtered***
- Sites where status is not "Proposed" or "Not Reported".
- Sites labelled as "Ramsar Site, Wetland of International Importance", with Designation Type "International".
- Sites designated under:
  - Ramsar Sites Criteria 4 (Sites that support plant and/or animal species at a critical stage in their life cycles, or provides refuge during adverse conditions);
  - Ramsar Sites Criteria 7 (Sites that support a significant proportion of indigenous fish subspecies, species or families, life-history stages, species interactions and/or populations that are representative of wetland benefits and/or values and thereby contributes to global biological diversity);
  - Ramsar Sites Criteria 8 (Sites that are an important source of food for fishes, spawning ground, nursery and/or migration path on which fish stocks, either within the wetland or elsewhere, depend); and
  - Ramsar Sites Criteria 9 (Sites that regularly support 1% of the individuals in a population of one species or subspecies of wetland-dependent nonavian animal species).
- All polygon data and point data with present, non-zero reported area (`REP_AREA > 0`).

***How the data are processed***
- Invalid S2 geometries (see Section 1) are made valid.
- Point data buffered to reported area, `REP_AREA`.
- Data are unioned to remove overlap (see Section 2).

##### 24.6 Protected areas under IUCN management categories I and II

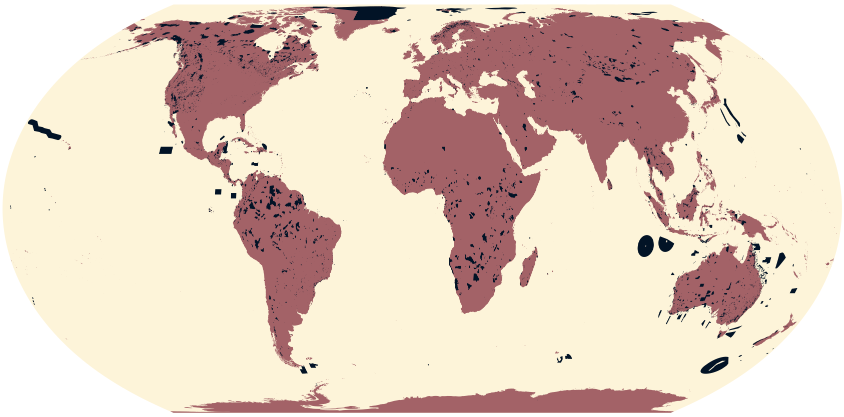

**Figure 52.** Global distribution of all protected areas under IUCN management categories I and II.

- **Number of records**: 33,720
- **Area**: 10,870,000 $km^2$
- **Critical habitat type**: Likely
- **IFC PS6 criteria triggered**: C4 (Highly Threatened or Unique Ecosystems)

***Justification and alignment with IFC PS6 criteria***
See Martin *et al.* 2015[2] and Brauneder *et al.* 2018[6]. Updated data.

***How the data are filtered***
- Sites where Status is not "Proposed" or "Not Reported".
- Sites where the reported IUCN Management Category is:
  - Ia (strict nature reserve);
  - Ib (wilderness area); or
  - II (national park).

***How the data are processed***
- Invalid S2 geometries (see Section 1) are made valid.
- Point data buffered to reported area, `REP_AREA`.
- Data are unioned to remove overlap (see Section 2).

#### 24.7 Natural and mixed World Heritage Sites

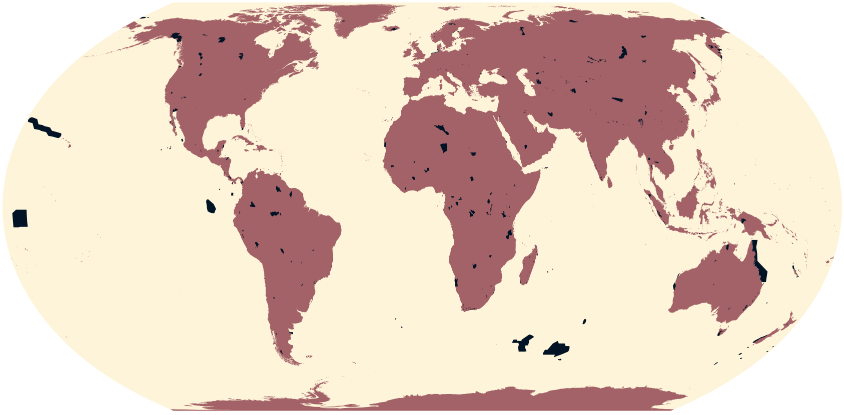

**Figure 53.** Global distribution of natural and mixed World Heritage Sites.

- **Number of records**: 266
- **Area**: 3,771,000 km$^2$
- **Critical habitat type**: Likely
- **IFC PS6 criteria triggered**: C4 (Highly Threatened or Unique Ecosystems)

***Justification and alignment with IFC PS6 criteria***

See Martin *et al.* 2015[2] and Brauneder *et al.* 2018[6]. Updated data.

***How the data are filtered***

- Sites where Status is not "Proposed", "Established", or "Not Reported".
- Sites labelled as "World Heritage Site (natural or mixed)", with Designation Type "International".

***How the data are processed***

- Invalid S2 geometries (see Section 1) are made valid.
- Point data buffered to reported area, `REP_AREA`.
- Data are unioned to remove overlap (see Section 2).
